# Probing DNA-protein interactions using single-molecule diffusivity contrast

**DOI:** 10.1101/2021.05.25.444860

**Authors:** Hugh Wilson, Miles Lee, Quan Wang

## Abstract

Single-molecule fluorescence investigations of protein-nucleic acid interactions require robust means to identify the binding state of individual substrate molecules in real time. Here we show that diffusivity contrast, widely used in fluorescence correlation spectroscopy at the ensemble level and in single-particle tracking on individual (but slowly diffusing) species, can be used as a general readout to determine the binding state of single DNA molecules with unlabeled proteins in solution. We first describe the technical basis of drift-free single-molecule diffusivity measurements in an Anti-Brownian ELetrokinetic (ABEL) trap. We then cross-validate our method with protein-induced fluorescence enhancement (PIFE), a popular technique to detect protein binding on nucleic acid substrates with single-molecule sensitivity. We extend an existing hydrodynamic modeling framework to link measured diffusivity to particular DNA-protein structures and obtain good agreement between the measured and predicted diffusivity values. Finally, we show that combining diffusivity contrast with PIFE allows simultaneous mapping of binding stoichiometry and location on individual DNA-protein complexes, potentially enhancing single-molecule views of relevant biophysical processes.

## Introduction

Single-molecule fluorescence measurements of protein-nucleic acid interactions^1–6^ have revealed new insight into the maintenance and processing of genomic information. In these experiments, access to the binding state of individual substrate molecules is essential and can be measured by labeling both the substrate and the protein with different reporters to look for colocalization^6,7^ or a Förster resonance energy transfer (FRET)^8,9^ signal. However, the requirement to fluorescently-tag the protein, often at site-specific locations, complicates the experimental design and limits the maximum allowable protein concentration due to background fluorescence^10,11^ (e.g. ~10 nM for total internal reflection microscopy).

Recently, protein-induced fluorescence enhancement (PIFE) has been developed as a powerful alternative to detect protein-nucleic acid binding without labeling the protein^5,12,13^. In the PIFE assay, a fluorescent probe on the DNA (or RNA) becomes brighter when an unlabeled protein binds in its vicinity and the degree of fluorescence enhancement depends on the dye-protein distance within a range of ~0-3 nm. This phenomenon is generally understood as leveraging the unique photophysical properties of cyanine dyes (e.g. Cy3, Cy5, DY547): the binding of a nearby protein suppresses the otherwise efficient, non-radiative photo-isomerization pathway from the dye’s singlet excited state, likely due to steric hindrance^14^. Since PIFE elegantly bypasses the requirement for protein labeling, it has gained popularity in recent years in many single-molecule studies of protein-nucleic acid systems^5,15–19^.

In this work, we aim to develop an alternative contrast mechanism to probe protein-nucleic acid interactions, using the physical properties of single molecules, without protein labeling. Measurements of hydrodynamic properties report on the global size and shape of macromolecules and are widely used to characterize biomolecular interactions at the ensemble^20,21^ and sub-ensemble^22^ levels. In addition, single-particle tracking^23^ is frequently used to detect interactions of individual biomolecules when diffusion is slow (*D* < 10 μm^2^/s), for example in membrane-bound^24^ or viscous cellular contexts^25,26^. Previously we developed a diffusometry platform based on an Anti-Brownian ELectrokinetic (ABEL) trap^27^ (Methods) which is capable of precisely measuring the translational diffusion coefficient (*D*) of individual, fast diffusing biomolecules (*D* ~ 100 μm^2^/s) in solution. Leveraging this advance, we reasoned that *D* would be a direct physical parameter to sense protein-nucleic acid interactions at the single-molecule level. In this work, we demonstrate this modality and cross-validate with PIFE. To link measured diffusivity values to particular molecular complexes, we extend the “HullRad” framework^28^ to model the diffusion coefficient of DNA-protein complexes which could be present in experiments. Further, we demonstrate the unique capability to independently resolve protein binding stoichiometry and binding location on a single short DNA molecule by combining diffusivity contrast with PIFE.

## Materials and Methods

### Sample preparation

DNA samples were ordered from Integrated DNA technology (IDT, Coralville, IA) and used without further purification. Detailed construct information including sequences and dye labeling positions can be found in Table S1. Duplexes were formed by mixing the labeled and the complimentary strands with a ratio of 1:1.5 in HN100 buffer (20 mM HEPES, pH8, 100 mM NaCl), heating to 90 °C for 2 min and gradually cooling to room temperature. Restriction enzymes (*BamHI* and *Eco*RI) were purchased from New England BioLabs (*BamHI:* R0136M, *EcoRI:* R0101M) and used without further purification. The stock concentrations of the enzymes (*Bam*HI: 1.1 μM, *Eco*RI: 0.74 μM) were provided by New England BioLabs. ABEL trap experiments were performed in a buffer containing 20 mM HEPES pH 7.8, 25 mM NaCl, 2 mM CaCl_2_ with 5-10 pM labeled DNA duplex, 1-10 nM *Bam*HI and/or 0.4-5 nM *Eco*RI. An oxygen scavenger system [50 or 100 nM protocatechuate 3,4-dioxygenase (OYC Americas) and 2 mM protocatechuic acid (Sigma)] and 2 mM Trolox were added to suppress Cy5 blinking and photobleaching^29^. The final sample solution also contained ~0.5 pM Atto647N-labeled ssDNA (5’-Atto647N-AAC TTG ACC C) which served as a fiducial marker of diffusion coefficient measurement consistency across different experimental runs.

### ABEL trap implementation

The ABEL trap was implemented similarly to previously published designs^27,30^ using an RM21 microscope frame (Mad City Labs). Briefly, we implemented a rapid, acousto-optic laser scanning and photon-by-photon mapping scheme to detect the position of a fast diffusing single molecule in solution. The detected molecule positions were refined by a hardware-coded (NI, PCIe7852R) Kalman filter^31^ before being used to compute the feedback voltages for molecule trapping. With minimum delay (~μs), these voltages were amplified and applied to a microfluidic sample chamber to counteract Brownian motion in two dimensions (x and y). The motion of the molecule in the longitudinal direction (z) was restricted by the depth of the chamber (~700 nm). In this work, excitation was provided by a 638 nm laser (Coherent Obis). The scanning pattern was chosen to be a 32-point “knight’s tour”^30^ (dwell time: 600 ns per point) which covers an area of approximately 3 μm×3 μm at the sample plane.

The microfluidic sample chamber was custom made in Princeton University’s nanofabrication facility on UV-grade fused silica wafers and passivated with polyethylene glycol (PEG)^32^. Specifically, following Piranha cleaning (3:1 mixture of sulfuric acid and hydrogen peroxide) and ~3 min sonication (or ~15 min incubation) in 1 M potassium hydroxide, the trapping chambers were incubated in a solution containing 500 mg mPEG-silane (MW:5 kDa, Laysan Bio), 10 ml ethanol, 250 μL water and 50 μL acetic acid (10 mM). The PEGylation reaction was allowed to proceed at room temperature for at least 48 hours in the dark. Before the experiment, the chamber was pre-treated with 0.5% Tween 20 in HN100 buffer (pH7.5)^33^ and rinsed extensively with ultrapure water.

### Single-molecule diffusometry

The diffusion coefficient (*D*) of a trapped single molecule was estimated using a maximum likelihood approach developed previously^27^. Briefly, the residual motion of a single molecule in the trap consists of (uncompensated) diffusion and voltage-induced electrokinetic drifts. Analysis of the motion trajectory with a recorded history of voltage inputs yields single-molecule motion parameters (both the diffusion coefficient *D* and the electrokinetic mobility). Because the photon-by-photon position tracking entails a high degree of localization uncertainty, an expectation-maximization framework was developed to iteratively achieve accurate position trajectory reconstruction and parameter estimation.

### Data analysis

All data analysis was performed using customized software written in Matlab. First, the raw arrival times of fluorescence photons were binned at 5 ms to identify regions that correspond to individual molecules. An intensity change point algorithm^34^ was applied to aid data segmentation. We used the following rules to identify the start and end times of a single molecule: (1) a brightness change from background to a stable high level is the beginning of a single-molecule event; (2) a brightness change from a stable high level to background is the termination of a single-molecule event; (3) a transient brightness spike is the signature of a second molecule diffusing into the trap and data immediately following the spike is assigned to a new molecule. After segmentation, brightness and diffusivity values associated with a single molecule were averaged and processed further.

## Results

### A focus lock improves single-molecule diffusometry stability

To differentiate molecular complexes by their hydrodynamic size, it is important to measure single-molecule diffusion coefficient (*D*) precisely and reproducibly. We accomplish this by using an ABEL trap to capture the molecule of interest in solution, allowing its diffusive motion to be continuously observed over many seconds. From statistical analysis of the residual in-trap motion, *D* of a single-molecule can be extracted with high precision^27^.We have previously shown that uncertainty in *D* measured with the ABEL trap scales with 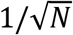 (N, the number of photons detected from the single molecule, Fig. S2). However, focus drift during recording can prevent the photon-limited precision from being reached, because defocus leads to a larger scanning beam at the trapping plane, which deteriorates the accuracy of the *D* estimation algorithm, Fig. 2C. To counteract focus drift and thus improve *D* estimation, we implemented a focus lock system similar to many other microscopy platforms^35–38^. In this approach, an auxiliary laser is reflected off the coverslip-sample interface near total internal reflection conditions and imaged onto an sCMOS camera (Fig. 2A). A change of objective-sample distance (*d*) results in a lateral translation of the beam (Fig. S1). The focus can be locked by commanding the z-piezo stage in a feedback loop to minimize the difference between the measured and target beam position (dashed horizontal line, Fig. 2A). Our system achieved a focus stability of ~8 nm (standard deviation, Fig. 2B) and effectively eliminated defocus-induced drift in diffusion estimation (Fig. 2D). With the focus locked, we achieved near photon-limited precision in *D* estimation over hour-long experiments (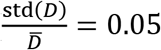, for *D*~50 μm^2^/s with one second measurement time, Fig. S2).

**Figure 1.**
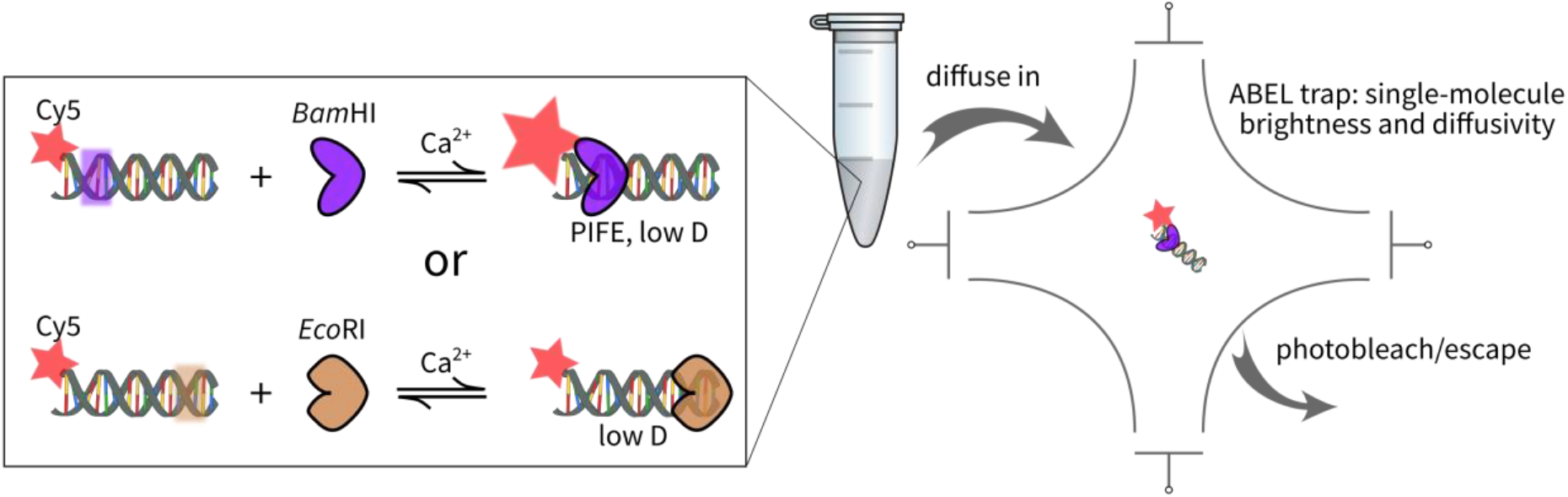
Probing DNA-protein interactions by single-molecule diffusivity contrast. The binding between short duplex DNA molecules (15-52bp in length) and two restriction enzymes is used to validate the method (left box). For each DNA length there is a *BamHI* binding site (purple shaded box) 2bp away from the terminal Cy5 label. Binding of *Bam*HI is expected to simultaneously enhance the brightness of Cy5 (via protein-induced fluorescence enhancement, PIFE) and lower the diffusion coefficient. Some DNA molecules also have an *Eco*RI binding site (brown shaded box) far from the Cy5 label and only a diffusivity change, not PIFE is expected upon *EcoRI* binding. The experiments are conducted by loading a sample containing labeled DNA (~pM concentration) and restriction enzyme (~nM concentration) into an ABEL trap microfluidic device (right schematic). Individual molecules are captured by feedback electrokinetic trapping and their brightness and diffusivity are determined.

**Figure 2.**
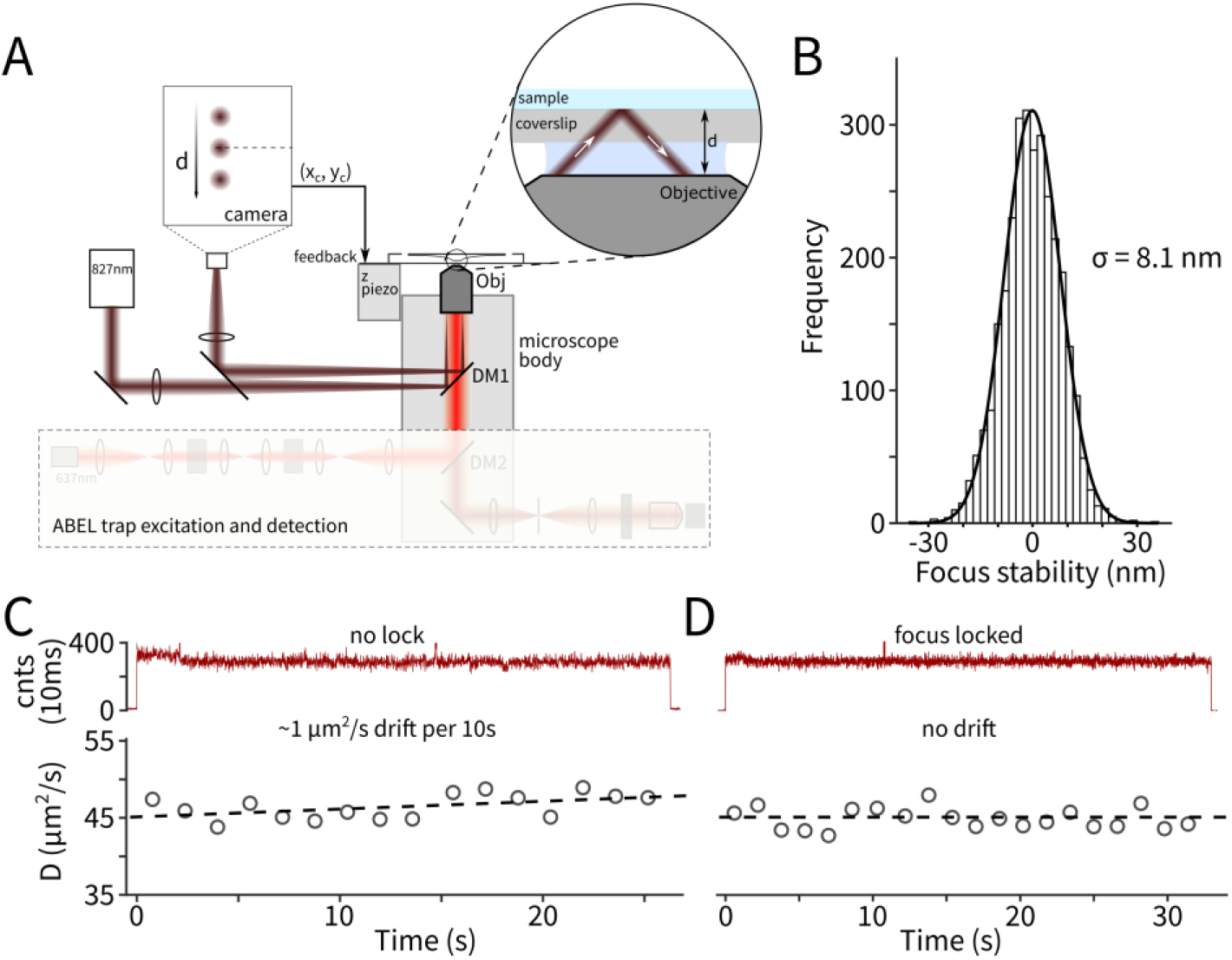
A focus lock system improves single-molecule D estimation. (**A**) Simplified schematic for the focus lock design. A near IR laser is coupled to the microscope in near total internal reflection mode and the reflected spot is imaged on to a camera. Changes in the objective-sample distance (d) result in lateral position (xc and yc) shifts of the camera spot. The difference between the spot position and a user-provided setpoint is used as the error signal for focus stabilization. (**B**) A representative distribution of deviations of the objective-sample distance (d) from the setpoint when the focus lock is engaged. The standard deviation (σ) is extracted by a Gaussian fit. (**C**) An example trapping trace recorded with severe focus drift. Top: brightness in 10ms bins. Bottom: estimated diffusion coefficient every 1.4 seconds. The apparent increase of diffusion coefficient is an artifact due to the excitation beam drifting out of focus. The dashed line is a linear fit of the data. (**D**) A representative trace recorded with the focus locked. The measured diffusion coefficient remains stable over time.

### Diffusion contrast maps single-molecule DNA-protein interactions with and without PIFE

To demonstrate the application of diffusion contrast measurements to DNA-protein interactions, we first designed an experiment to cross-validate our approach with PIFE. We measured a 15bp duplex DNA construct with a *Bam*HI binding site (GGATCC) (Table S1) with and without BamHI in solution. With Ca^2+^ in place of Mg^2+^ in our buffer solution, *Bam*HI is expected to bind stably at the recognition sequence without inducing DNA cleavage^39^. The DNA substrate is fluorescently labeled with a Cy5 dye at the terminal base and the separation between Cy5 and the binding site is 2 base pairs. Previously it was demonstrated that at such a small distance, Cy5 displays a robust PIFE response^14^. As a result, in our experiment, specific binding of *Bam*HI is expected to lower the diffusion coefficient of the DNA and simultaneously increase Cy5 brightness.

Without *Bam*HI, individual DNA duplexes (Fig. 3A top) showed homogenous brightness (~110 cnts/5ms) and diffusion coefficient (~115 μm^2^/s) (Fig. 3A bottom). When the measurement was conducted in the presence of 10 nM *Bam*HI (with Ca^2+^ in place of Mg^2+^ to prevent DNA cleavage), two different populations were seen (Fig. 3B). One population was evidently the bare DNA, as its brightness and diffusivity values closely resembled those observed in the DNA only measurement. The other population displayed the behavior expected from specifically bound *Bam*HI-DNA complexes: a higher brightness (~182 cnts/5ms) due to PIFE and a lower diffusion coefficient (~68 μm^2^/s). Further, this high-brightness, low-D population decreased in abundance when a ten-fold lower concentration (1 nM) of *Bam*HI was used (Fig. S3), and was not observed in control experiments with mutant DNA (containing a single base change in the recognition sequence, Fig. S4A) or a non-cognate protein (*EcoRI,* Fig. S4B). We observed a fluorescence enhancement factor of 1.65 fold, consistent with previous measurement^12^ (1.9 fold enhancement at 1bp away). In addition, the measured diffusion coefficient of the *Bam*HI-DNA complex (~68 μm^2^/s) agrees with structure guided modeling (see the section below: *“Modeling the diffusion coefficients of DNA-protein complexes using extHullRad”*). Importantly, all molecules identified as *Bam*HI-DNA complexes by diffusivity contrast displayed enhanced brightness, demonstrating excellent consistency of both modalities in this example. Nevertheless, a small fraction of the *Bam*HI-bound complex showed lower fluorescence enhancement. We speculate that this could be due to heterogeneity in Cy5’s photophysical behavior or *Bam*HI’s binding geometry.

**Figure 3.**
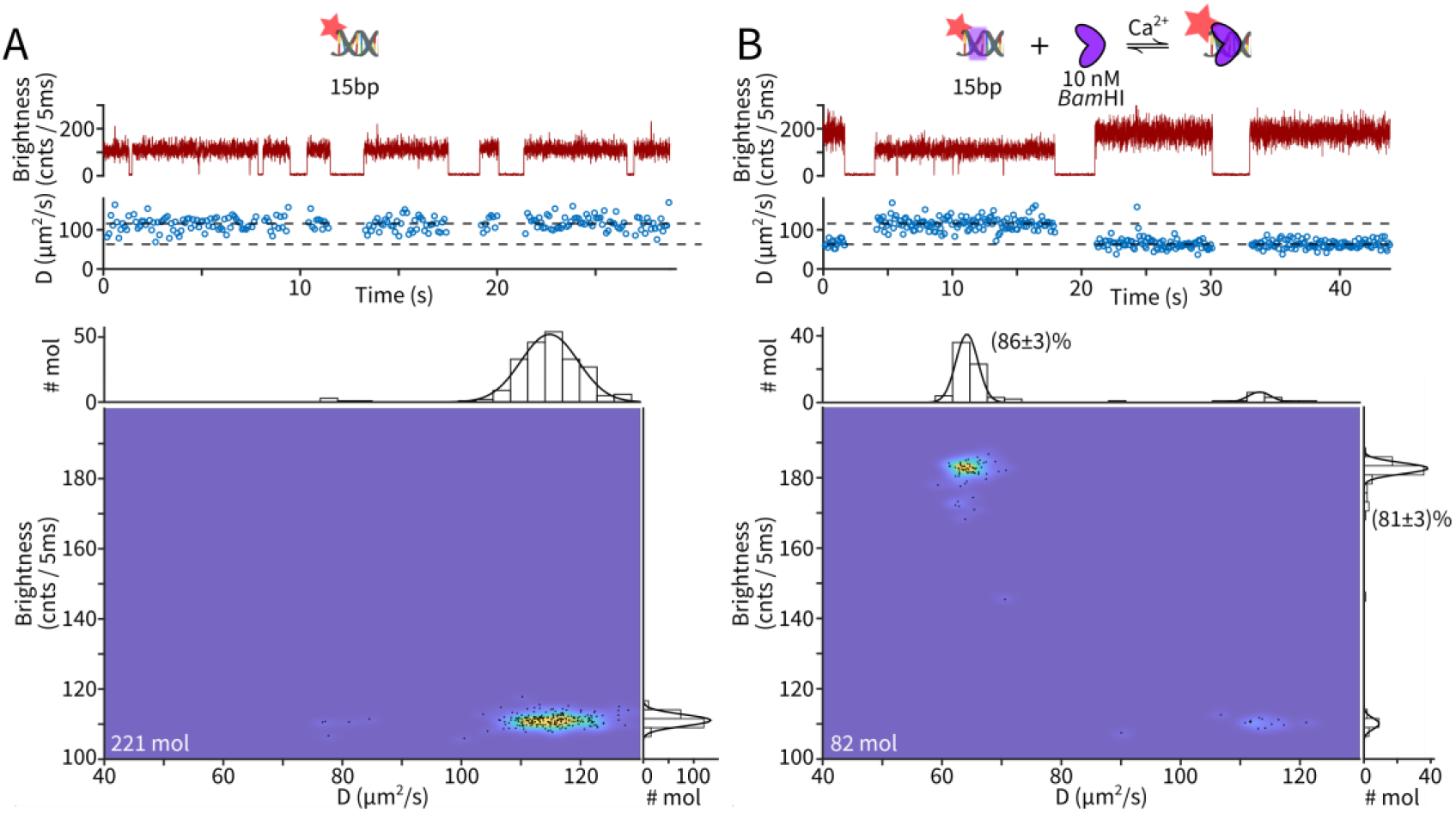
Probing *BamHI* binding to 15bp duplex DNA using PIFE and diffusivity contrast. (**A**) DNA without *Bam*HI. Top: representative time trace showing single-molecule brightness (5 ms bins) and measured diffusivity (every 100 ms) Bottom: D-Brightness scatter plot of 221 measured molecules. Every black dot represents the diffusivity and brightness of a single molecule averaged over the trapping period. The underlying density is estimated using a 2D kernel density estimator. Marginal histograms are fit with a single-component Gaussian function. (**B**) DNA with 10 nM *Bam*HI. Top: representitave time trace showing single-molecule brightness (5 ms bins) and measured diffusivity (every 100 ms). Bottom: D-Brightness scatter plot of 82 molecules. Both marginal histograms are fit with a two-component Gaussian function and the relative abundance of the major population is extracted from the fit.

We next measured DNA-protein binding without PIFE. We used a Cy5-labeled, 40bp duplex DNA with an *Eco*RI binding site (GAATTC) 27 base pairs away from the terminal Cy5. Previous work suggested that PIFE should not be detectable at this separation^12^. Indeed, when we probed the DNA substrates molecule-by-molecule in the presence of 1 nM *Eco*RI, we observed homogeneous brightness among the molecules probed (Fig. 4 top). The diffusion coefficient, on the other hand, displayed two distinct levels that correspond to bare and *Eco*RI-bound DNA (Fig. 4), respectively. (A minor population with D~95 μm^2^/s and brightness ~110 cnts/5ms was also observed and identified as un-hybridized, Cy5-labeled single-strand DNA.) Note that in this example, conventional single-molecule fluorescence microscopy, which records only fluorescence brightness, would not be able to differentiate bound and unbound DNA molecules. Diffusivity contrast, on the other hand, provides a general and reliable mapping of DNA-protein interactions.

**Figure 4.**
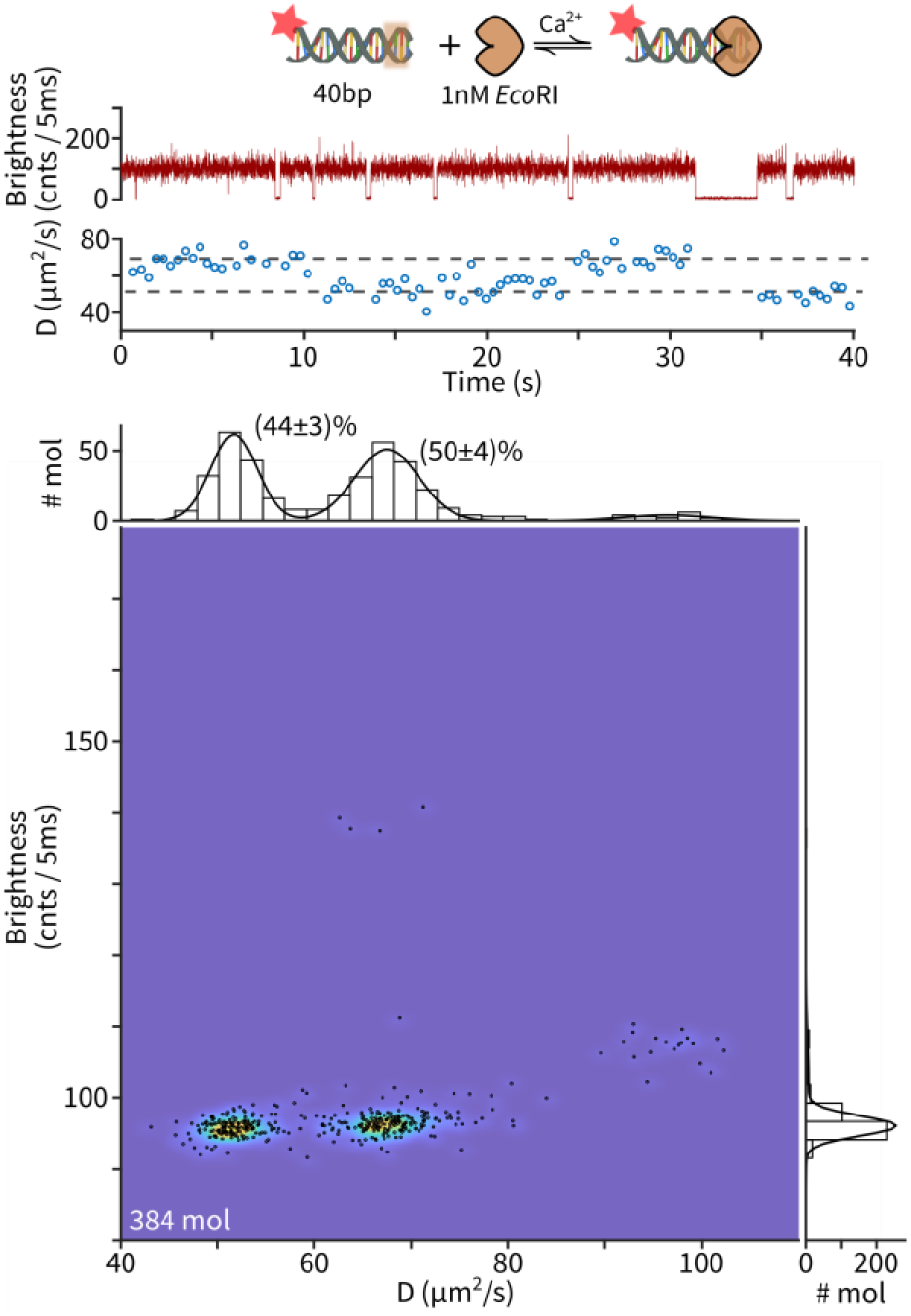
Diffusivity contrast maps *Fco*RI-DNA interactions without PIFE. Top: representative time trace showing single-molecule brightness (5 ms bins) and measured diffusivity (every 100 ms) Bottom: D-Brightness scatter plot of 384 measured molecules. Every black dot represents the diffusivity and brightness of a single molecule averaged over the trapping period. The underlying density is estimated using a 2D kernel density estimator. The marginal histogram along the brightness axis is fit with a single-component Gaussian function while the marginal histogram along the diffusivity axis is fit with a three-component Gaussian function. The relative abundance of the two major populations are extracted from the fit.

### Modeling the diffusion coefficient of DNA-protein complexes using extHullRad

We next developed a modeling pipeline to associate measured diffusivity values with particular DNA-protein complexes. Predicting the diffusion coefficient of biological macromolecules from atomic coordinates is well-established^40^ and many accessible and easy-to-use tools^28,41^ are available. However, these modeling tools cannot be directly applied to the samples in our experiments due to the lack of atomic structures for our full DNA-protein complexes (available database structures of DNA-binding proteins are often determined with very short or no substrate DNA). Inspired by the recently developed HullRad framework^28^, we recognize that without full structural data, the hydrodynamic properties of DNA-protein assemblies can still be modeled by a composite approach based on constructing the smallest envelope (convex hull) that encompasses the entire complex in 3D. Our pipeline (“extHullrad”, Fig. 5A) starts with an existing protein-DNA structure. The DNA portion is replaced with a cylindrical rod (radius of 1 nm and 0.34 nm rise per base pair), aligned to the DNA axis and binding site in the original structure. This allows duplexes of different lengths (much shorter than the 40-50 nm persistence length of dsDNA^42,43^) to be modeled without molecular details. The composite structure is then used as the input to the original HullRad algorithm^28^(Fig. 5A inner square), in which the convex hull and hydration shell around a molecular complex is approximated as a prolate ellipsoid, and an extension of the Stokes-Einstein equation is used to calculate the translational diffusion coefficient (see ref. 28 for details). The extHullRad routine is implemented in Python and available as supplementary code.

**Figure 5.**
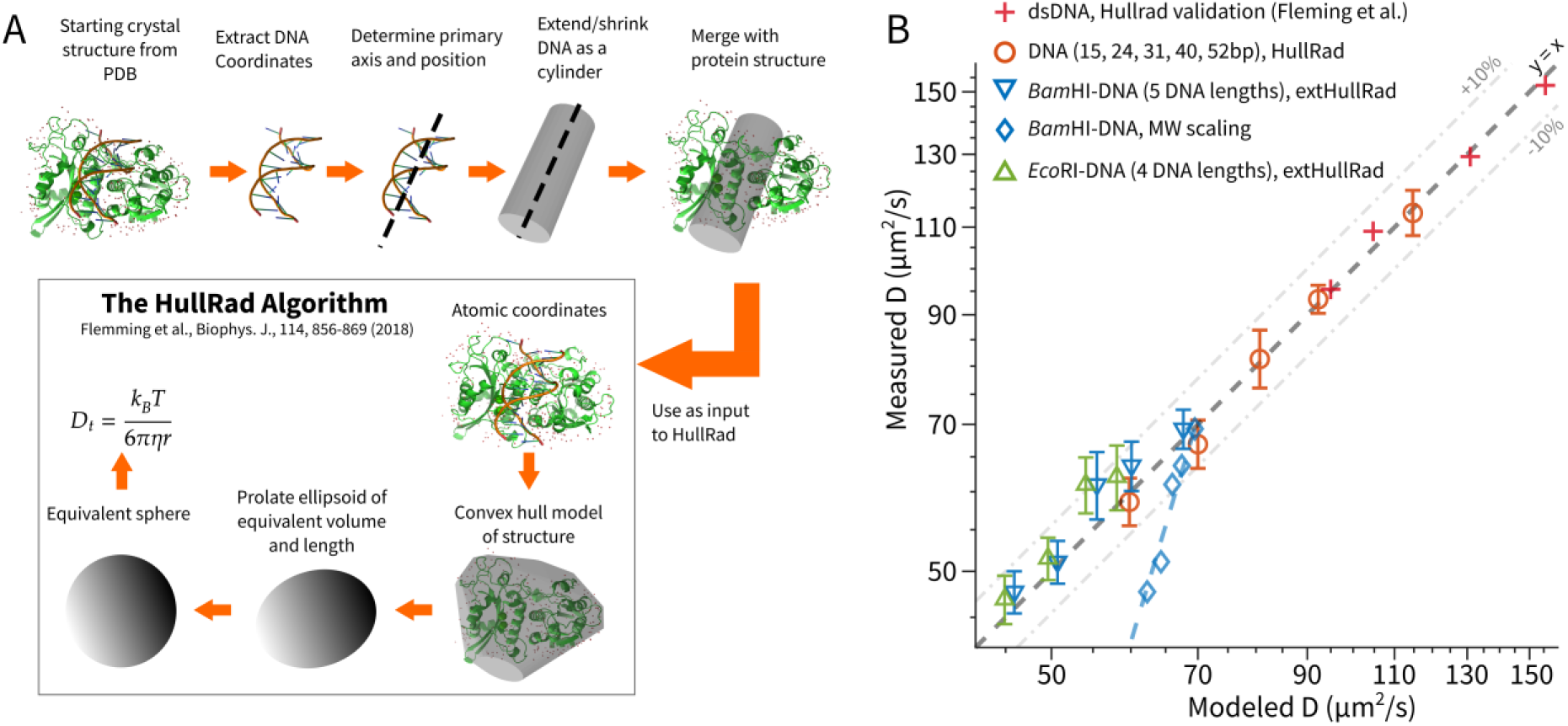
extHullRad successfully predicts the diffusion coefficient of DNA-protein complexes. (**A**) Workflow of extHullRad (see text for details) (**B**) Comparison of modeled and measured diffusivity values across different samples. Red plus signs: short duplex DNA molecules (reproduced from Fleming et al.) which were used to validate the modeling accuracy of the original HullRad framework. Orange circles: DNA molecules from this study (15, 24, 31, 40 and 52bp) measured using focus-stabilized single-molecule diffusometry and modeled using HullRad. Blue triangles: *Bam*HI-bound 15, 24, 31, 40 and 52bp dsDNA, measured in this work and modeled using extHullRad. Green triangles: *Eco*RI-bound 24, 31, 40, 52bp dsDNA, measured in this work and modeled using extHullRad. Blue diamonds: comparison of the measured D of *Bam*HI-DNA complexes with a naïve molecular weight scaling D prediction model (D ∝ MW^-1/3^). Error bars of measured *D* represent standard deviation values from Gaussian fits of *D* histograms (Figs. S5-S9).

In Fig. 5B we compare the modeling results to single-molecule diffusometry measurements. We first confirm that measured D values of bare DNA (15, 24, 31, 40 and 52bp) agree with Hullrad modeling (Fig. 5B, orange circles). We then compare measurements of two different proteins (*Bam*HI and *Eco*RI) bound to substrate dsDNA of different lengths (15-52bp, nine datasets in total, Figs. S5-S9). For each condition, we use the exHullRad pipeline to model the diffusion coefficient of the DNA-protein complex (starting from PDB:2BAM for *Bam*HI, and 1ERI for *Eco*RI) and compare to experimental results (Figs. S5-S9). We achieved satisfactory agreement between the modeled and measured D values of the complexes (a mean error of 5.2% across 9 complexes, Fig. 5B, triangles). Importantly, it is critical to model the *shape* of the DNA-protein complexes used here, a task that extHullRad was designed to accomplish. A naïve prediction which assumes complexes are spherical and uses *D* ∝ *MW*^-1/3^ scaling (MW = molecular weight) produced much larger discrepancies (Fig. 5B diamonds).

### Diffusion contrast with PIFE probes ternary complexes with sequence context

Finally, we demonstrate that combining diffusivity contrast and PIFE enables three-component DNA-protein reactions to be resolved at the single-molecule level. We used a 40bp DNA duplex with both *Bam*HI and *Eco*RI binding sites and aimed to identify all possible binding configurations in the presence of the two restriction enzymes (Fig. 6A). As above, the PIFE-sensitive dye Cy5 is placed 2bp away from the *Bam*HI site and 27bp away from the *Eco*RI site. We anticipated that PIFE would report the occupancy of the *Bam*HI site and diffusivity would probe the number of protein molecules bound (i.e. 0, 1, 2), thus giving rise to four distinct populations in the *D*-brightness space (Fig. 6B). Experiments at several concentrations of *Eco*RI confirmed this pattern with four populations clearly visible (Fig. 6C-D). The two populations at lower brightness (~95 cnts/5ms), one with *D* around 67 μm^2^/s and the other with *D* ~ 53 μm^2^/s, can be assigned as unbound DNA and DNA-*Eco*RI complex, respectively, based on the absence of PIFE and comparison to data shown in Fig. S8 (DNA-only and DNA with one restriction enzyme each). The two populations at higher brightness (160 cnts/5ms) have *Bam*HI bound. Among these, the *D* ~ 53 μm^2^/s population is the DNA-*Bam*HI binary complex based on a similar *D* to DNA-*Eco*RI and comparison to Fig. S8B. Finally, the population with the lowest *D* (~45 μm^2^/s) can be assigned to the fully assembled, ternary complex (DNA-*Bam*HI-*Eco*RI) based on the high brightness and the larger hydrodynamic size compared with DNA-*Bam*HI. Furthermore, the relative abundance of the populations shifted as the *Eco*RI concentration was varied from 0.4 nM to 2 nM (Fig. 6C-D and S10), in a way that is consistent with the expected *Eco*RI binding-site occupancy of each state in our assignment. Note that in this example, neither the PIFE signal nor the diffusion coefficient alone would be able to resolve all binding configurations.

**Figure 6.**
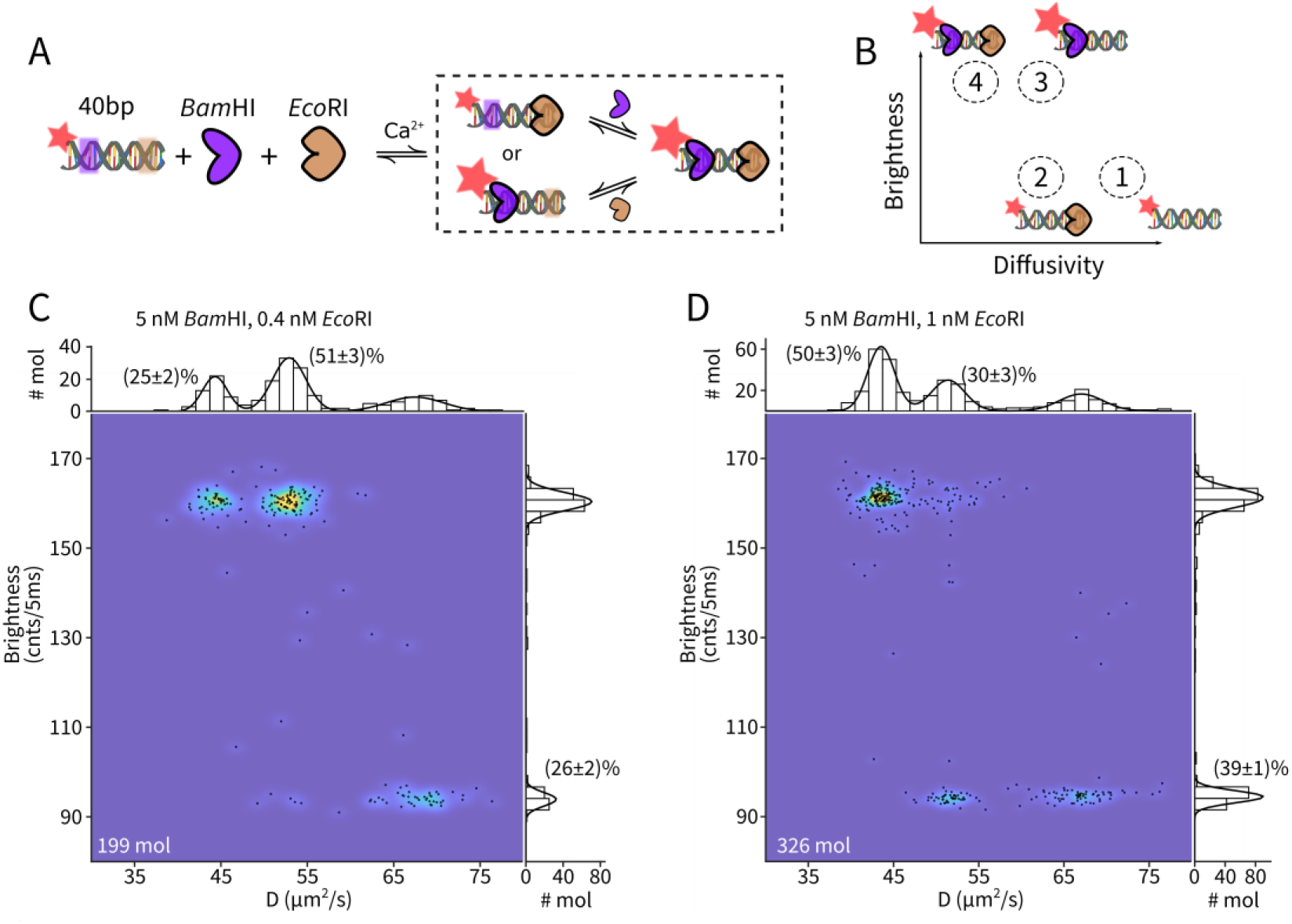
Combining single-molecule diffusivity contrast and PIFE identifies all four binding configurations between a dual binding-site DNA and two proteins. (**A**) Experimental design: a 40bp DNA containing both *Bam*HI and *Eco*RI binding-sites can undergo sequential binding to arrive at the DNA-*Bam*HI-*Eco*RI ternary complex. (**B**) Four possible binding configurations are possible in solution and can be resolved in the D-brightness parameter space. (**C**) Experimental D-Brightness scatter plot of 199 measured molecules with 5 nM *Bam*HI and 0.4 nM *Eco*RI. (**D**) Experimental D-Brightness scatter plot of 326 measured molecules with 5 nM *Bam*HI and 1 nM *Eco*RI. In panels C and D, every black dot represents the diffusivity and brightness of a single molecule averaged over the trapping period. The underlying density is estimated using a 2D kernel density estimator. The marginal histogram along the brightness axis is fit with a two-component Gaussian function while the marginal histogram along the diffusivity axis is fit with a three-component Gaussian function. The relative abundance of the populations are extracted from the respective fits.

## Discussion

This work establishes single-molecule diffusivity as a general readout to probe DNA-protein interactions in solution. In contrast to many classic biochemical approaches for analyzing protein-nucleic acid interactions (e.g. the electrophoretic mobility shift assay^44^, which separates molecular complexes into distinct gel bands), the method presented here operates on individual molecules in solution at chemical equilibrium. From analyzing the relative abundance of the protein-bound and free DNA species, we estimated a K_d_ of ~2-5 nM for *Bam*HI and ~1 nM for *Eco*RI under the buffer condition (20 mM HEPES pH 7.8 with 25 mM NaCl and 2 mM CaCl_2_) and sequence contexts used in this study. These values are in line with recent solution phase experiments using fluorescence anisotropy (K_d_ ~9 nM for *Bam*HI with Ca^2+^)^45^ but larger than values measured previously using a filter binding assay^39^ (K_d_ ~ 0.2 nM for *Bam*HI) and a mobility shift assay^46^ (K_d_ ~ 0.4 nM for *Eco*RI without Ca^2+^). The discrepancy may be due to differences in assay medium, sequence context or the influence of a fluorophore close to the binding site. Beyond an equilibrium measurement of binding affinity, observing single complexes under equilibrium also offers a window into the dynamics and heterogeneity of assembly (Fig. S11). Information on the kinetics and relative order of binding steps may shed new light on the favored pathways of protein-nucleic acid assembly.

At the single-molecule level, diffusivity contrast provides an alternative to PIFE for sensing the binding between labeled nucleic acid and unlabeled protein. Using a specially designed DNA substrate, we have verified that both methods reliably identify protein binding. Each modality has its advantages and disadvantages. PIFE is straightforward to implement on existing single-molecule fluorescence platforms but requires strategically placing a sensor dye within a few nanometers of the binding site. Furthermore only a limited selection of dyes (mostly cyanine) are PIFE sensitive. Diffusivity contrast requires sophisticated hardware and analysis algorithms to implement but relaxes design constraints associated with PIFE. There is no restriction on dye type or labeling location on the nucleic acid. Importantly, PIFE is a photophysical phenomenon and combining with other spectroscopic modalities (e.g. single-molecule FRET) often results in compounded signals requiring sophisticated analysis and experimental design to delineate^47,48^. Stacking of cyanine dyes at the end of DNA duplexes^49^ as well as protein-dye orientation^50^ can also complicate the interpretation of PIFE signals^51^. Diffusivity contrast, on the other hand, is based on a physical property of molecular complexes (i.e. hydrodynamic size) and is completely orthogonal to spectroscopic readouts. Combining diffusivity contrast with other spectroscopic readouts adds an extra dimension^32^ that enhances single-molecule measurements. Moreover, we showed that diffusivity contrast provides complex-level structural information through the extHullRad modeling framework. Meanwhile, the time resolution of PIFE (<10ms) is much higher than our current implementation of diffusivity measurement (~200 ms), making PIFE a more suitable method to probe rapid binding/unbinding events.

Given the complementary nature of PIFE and diffusivity contrast, combining the two modalities brings new capabilities. We have demonstrated simultaneous mapping of stoichiometry (“how many protein molecules are bound”) and sequence context (“where are they bound”) in the interactions between one DNA substrate and two proteins. Similar ideas can be used to differentiate specific and non-specific binding. We envision that these capabilities will enable new single-molecule measurements of many fascinating phenomena in protein-nucleic acid interactions, for example, target search^52^ and DNA allostery^53^.

## ACKNOWLEDGMENTS

We thank Haw Yang and his lab for feedback and suggestions, Patrick Fleming for discussion of the extHullRad framework, New England Biolabs customer support for providing concentration values of their *Bam*HI and *Eco*RI products. This work is funded by the Lewis-Sigler Fellowship to QW and US Department of Energy Office of Basic Energy Sciences and Photosynthetic Systems Grant DE-SC0002423.

## SUPPORTING MATERIAL

Supporting material includes one table, eleven supplementary figures and supplementary code.

## AUTHOR CONTRIBUTIONS

HW, ML and QW designed the research. HW and QW performed the experiments and analyzed the data. ML developed the extHullRad framework.

