## Supplementary Information for "Probing DNA-protein interactions using single-molecule diffusivity contrast"

**Table S1. Sequences of the DNA strands used in this work**

| Strand | Sequence |
|---|---|
| 15bp substrate with a *Bam*HI binding site | 5'-Cy5-TA**G GAT CC**A TAG TAG-3' |
| | 5'-CTA CTA T**GG ATC C**TA-3' |
| 15bp non-cognate for *Bam*HI (single mutation) | 5'-Cy5-TAG GTT CCA TAG TAG-3' |
| | 5'-CT ACT ATGGAA CCTA-3' |
| 24bp substrate with both *Bam*HI and *Eco*RI binding sites | 5'-Cy5-TA**G GAT CC**A TA**G AAT TC**A GCG TAG-3' |
| | 5'-CTA CGC T**GA ATT C**TA T**GG ATC C**TA-3' |
| 31bp substrate with both *Bam*HI and *Eco*RI binding sites | 5'-Cy5-TA**G GAT CC**A TAG TTA GCG **GAA TTC** AGC GTA G-3' |
| | 5'-CTA CGC T**GA ATT C**CG CTA ACT AT**G GAT CC**T A-3' |
| 40bp substrate with both *Bam*HI and *Eco*RI binding sites | 5'-Cy5-TA**G GAT CC**A TAG TGA CGT ACG GTA GCG **GAA TTC** AGC GTA G-3' |
| | 5'-CTA CGC T**GA ATT C**CG CTA CCG TAC GTC ACT AT**G GAT CC**T A-3' |
| 52bp substrate with both *Bam*HI and *Eco*RI binding sites | 5'-Cy5-TA**G GAT CC**A TAG TGA GAG TAG CAA GTG CGT ACG GTA GCG **GAA TTC** AGC GTA G-3' |
| | 5'-CTA CGC T**GA ATT C**CG CTA CCG TAC GCA CTT GCT ACT CTC ACT AT**G GAT CC**T A-3' |
| Fiducial D marker | 5'-Atto647N-AAC TTG ACC C |

**Underscore**: *Bam*HI binding site

**Red**: *Eco*RI binding site

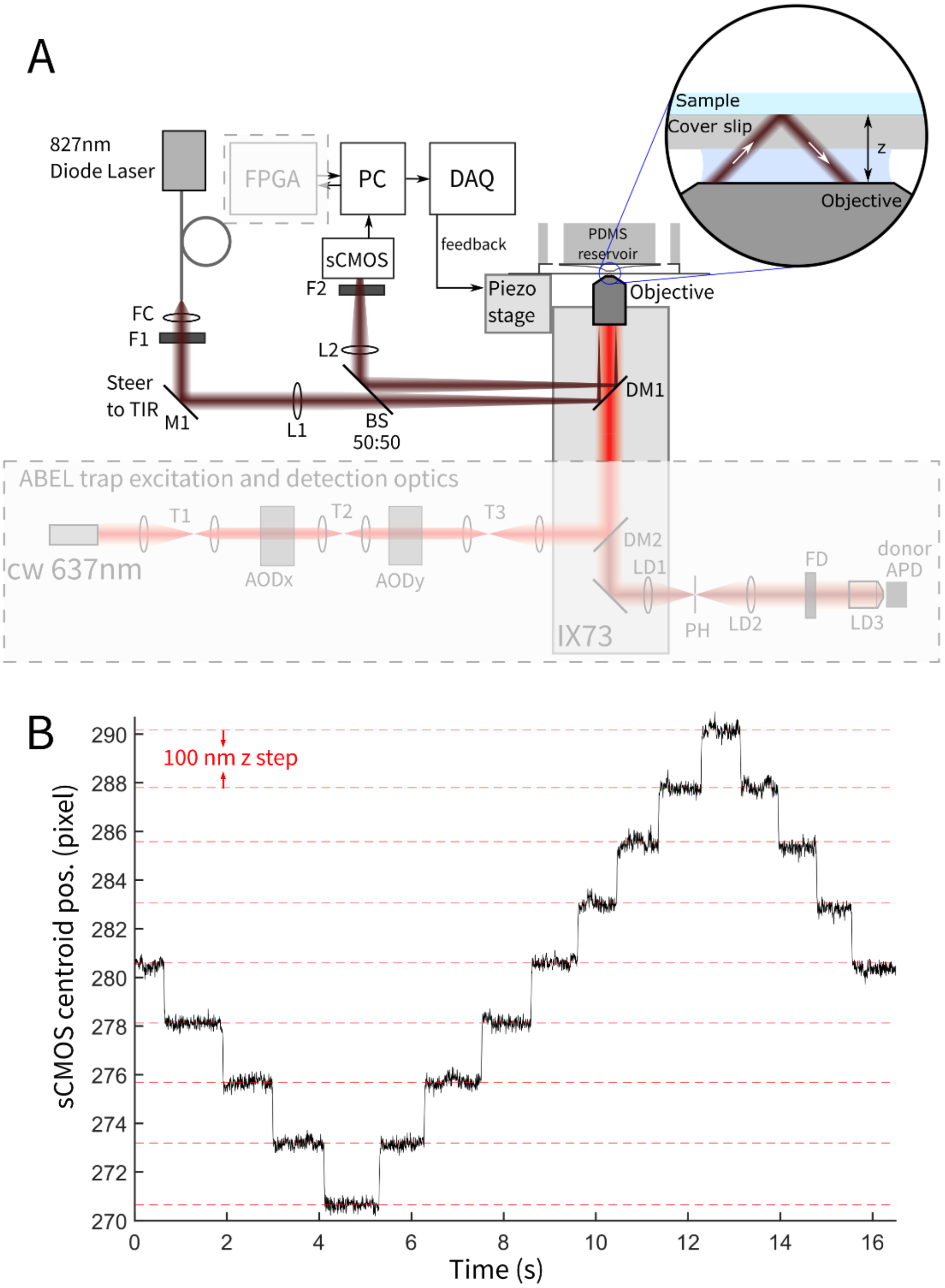


**Figure S1 Implementation of the focus lock system for improved single-molecule diffusometry.** (**A**) Schematic of the focus-lock design. 827nm Diode Laser: Thorlabs, LPS-830-FC, FC: fiber collimator (Thorlabs, F280FC-850), F1, F2: 800nm long pass filter (Thorlabs, FELH0800), M: mirrors (Thorlabs, BB1-EO3), L1 and L2: achromatic lenses with NIR coating (Thorlabs, LA1*-B, * specifies focal length, L1 = 250mm, L2 = 200mm), BS 50:50: beamsplitter

(Thorlabs, EBS1), DM1: 805nm short pass dichroic mirror (Thorlabs, DMSP805R). Objective: Silicone immersion objective (Olympus, 100X SI OIL NA1.35 UPLSAPO), z Piezo stage: Mad City Labs, Nano OP-65, sCMOS camera: PointGrey CM3-U3-31S4M-CS, PC: computer running Windows 7 and LabView 2016, DAQ: NI PCIe-6323, FPGA: NI PCIe-7852R. The ABEL trap excitation and detection optics were described in Ref.[1]. (**B**) Centroid position (pixels) of the focus tracking laser spot on the sCMOS camera, in response to 100nm steps of the piezo stage in open loop (focus lock off).

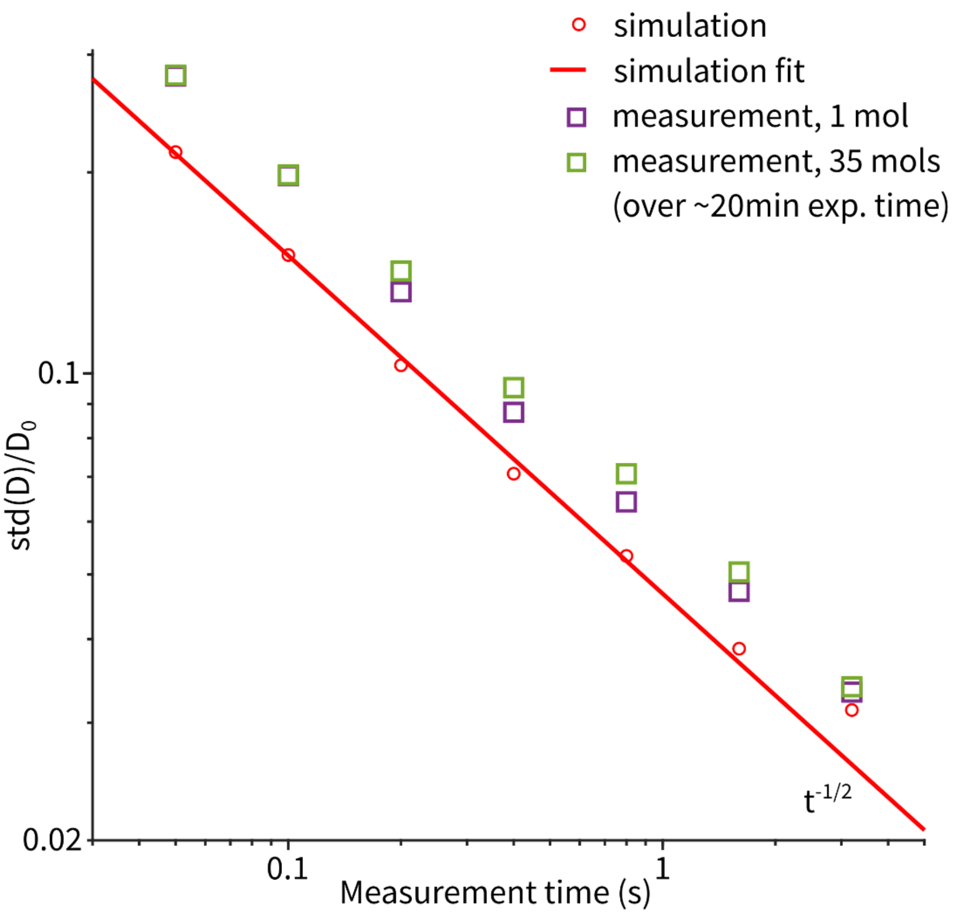


**Figure S2 Statistical uncertainty (standard deviation of D normalized by the mean) of single-molecule diffusivity as a function of measurement time.** Red circles: characterization using simulated ABEL trap data (see Ref. [2]). The following parameters, extracted from experiments, were used in the simulations. D = 49 $\mu m^2/s$, $\mu$ = 200 μm/s/V (electrokinetic mobility), w = 0.70 μm ($1/e^2$ beam radius of the scanning laser), bg = 1.09 kHz (background count rate), C = 20 kHz (averaged fluorescence count rate in the trap), $G_{fb}$ = 10 V/μm (feedback gain). To characterize the uncertainty of D estimation, the simulation was run 10 times, each with a total duration of 3.5 seconds. For each measurement time (T), the simulated traces were divided into consecutive, non-overlapping segments of duration T and the diffusion coefficient of each segment was estimated. The standard deviation of all estimated D values was computed and divided by the mean. The red line is a power-law fit to the simulated uncertainties as a function of measurement time. The fitted exponent is –(0.48 ± 0.02). Magenta squares: experimentally obtained relative uncertainty in D of one measured molecule. The sample is 52bp substrate DNA in the presence of 2nM *Eco*RI. The molecule characterized here (trapped for a total of 51.1 seconds) had a mean D value of 49.1 $\mu m^2/s$ and was classified as an *Eco*RI-52bp DNA complex. Similar to the characterization of the simulated data, the trace was divided into consecutive, non-overlapping segments of duration T and the D of each segment was estimated. The standard deviation of all estimated D values was computed and divided by the mean. Green squares: the same analysis performed on data pooled from 35 molecules measured over a 20-min span. For all measurements the focus lock was engaged. The precision obtained on 35 molecules is near-identical to the precision obtained on one molecule and both are close to the photon-limited bound (red line). These observations suggest that the measurement is stable over the 20min acquisition time with the focus lock.

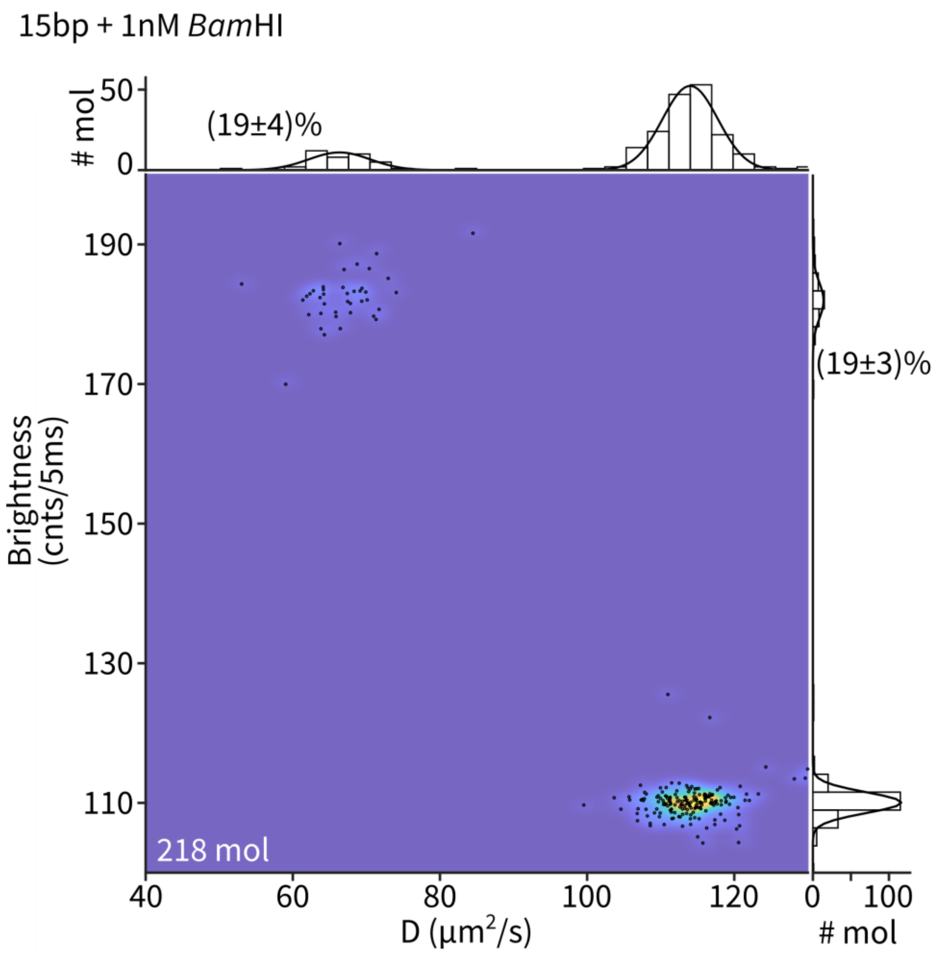


**Figure S3 Diffusivity-brightness mapping of single 15bp DNA molecules in the presence of 1 nM *Bam*HI.** Every black dot represents the diffusivity and brightness of a single molecule averaged over the trapping period. The underlying density is estimated using a 2D kernel density estimator. Marginal histograms are fit with a two-component Gaussian function. Under this condition, about 20% of dsDNA molecules were bound with a *Bam*HI protein.

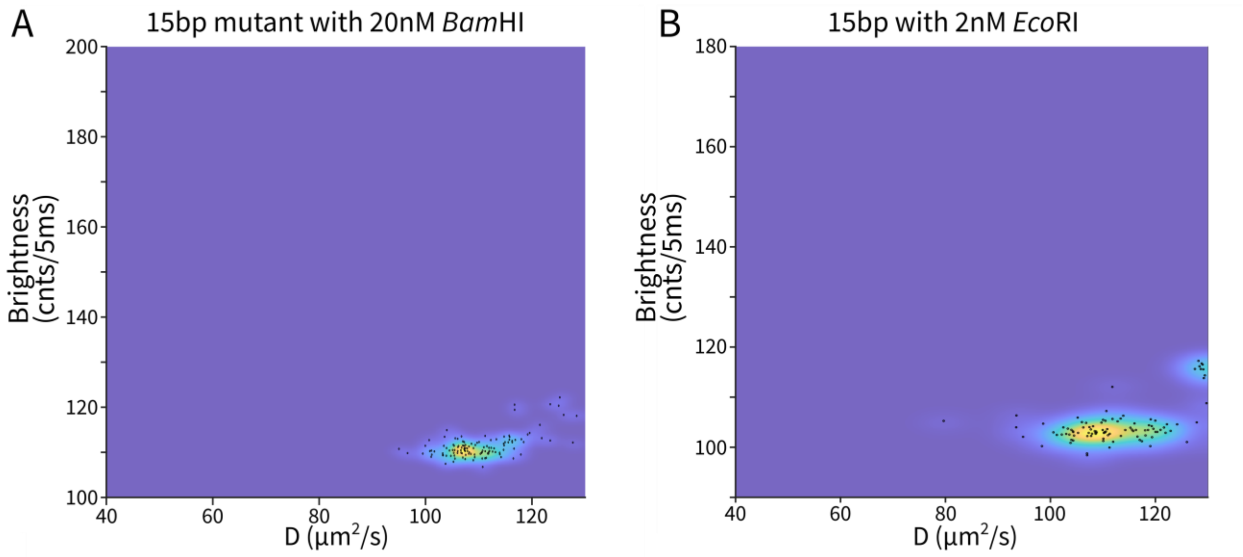


**Figure S4 Control experiments of 15bp DNA-*Bam*HI binding.** (**A**) Diffusivity-brightness scatter plot of 15bp-mut1 DNA with 20 nM *Bam*HI. Here the DNA substrate contains "GGTTCC" instead of the "GGATCC" recognition sequence. (**B**) Diffusivity-brightness scatter plot of 15bp DNA (*Bam*HI cognate) in the presence of 2 nM *Eco*RI. In both experiments, no stable protein binding was observed.

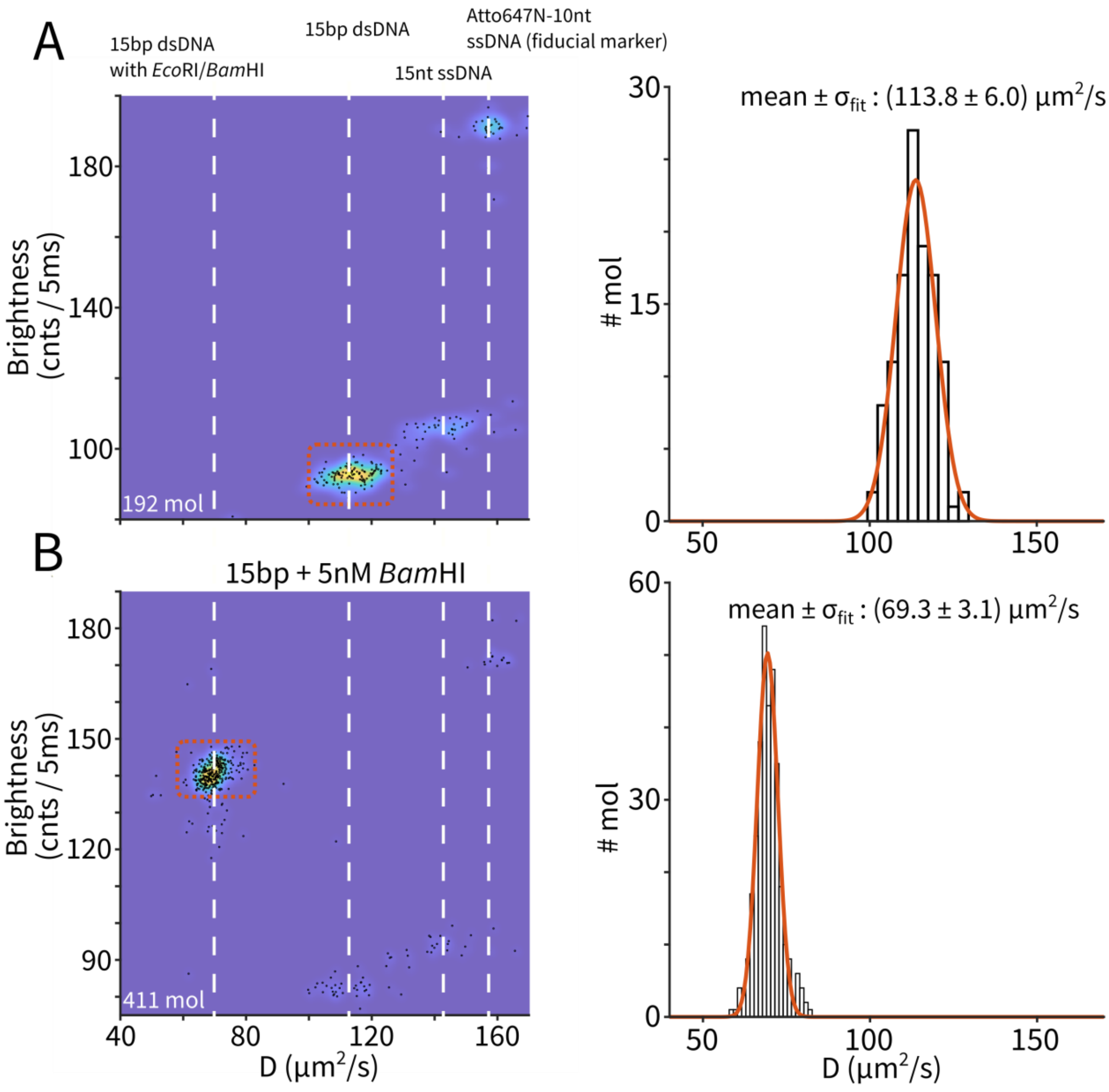


**Figure S5 Quantitative diffusivity measurement of 15bp DNA and 15bp-*Bam*HI complexes.** (**A**) Full diffusivity-brightness scatter plot of the 15bp DNA measurement. White dashed lines denote the molecular populations identified based on diffusion coefficient. Minor populations of unhybridized Cy5-ssDNA and the fiduciary marker DNA are present. The distribution of measured D values of the 15bp dsDNA only population (indicated by the orange box) is shown on the right. The mean and standard deviation of D were quantified by fitting the distribution with a Gaussian distribution (orange curve). (**B**) Full diffusivity-brightness scatter plot of the 15bp DNA measured with 5nM *Bam*HI. The distribution of measured D for the 15bp-dsDNA-*Bam*HI population (indicated by the orange box) is shown on the right. The mean and standard deviation of D were quantified by fitting the distribution with a Gaussian distribution (orange curve).

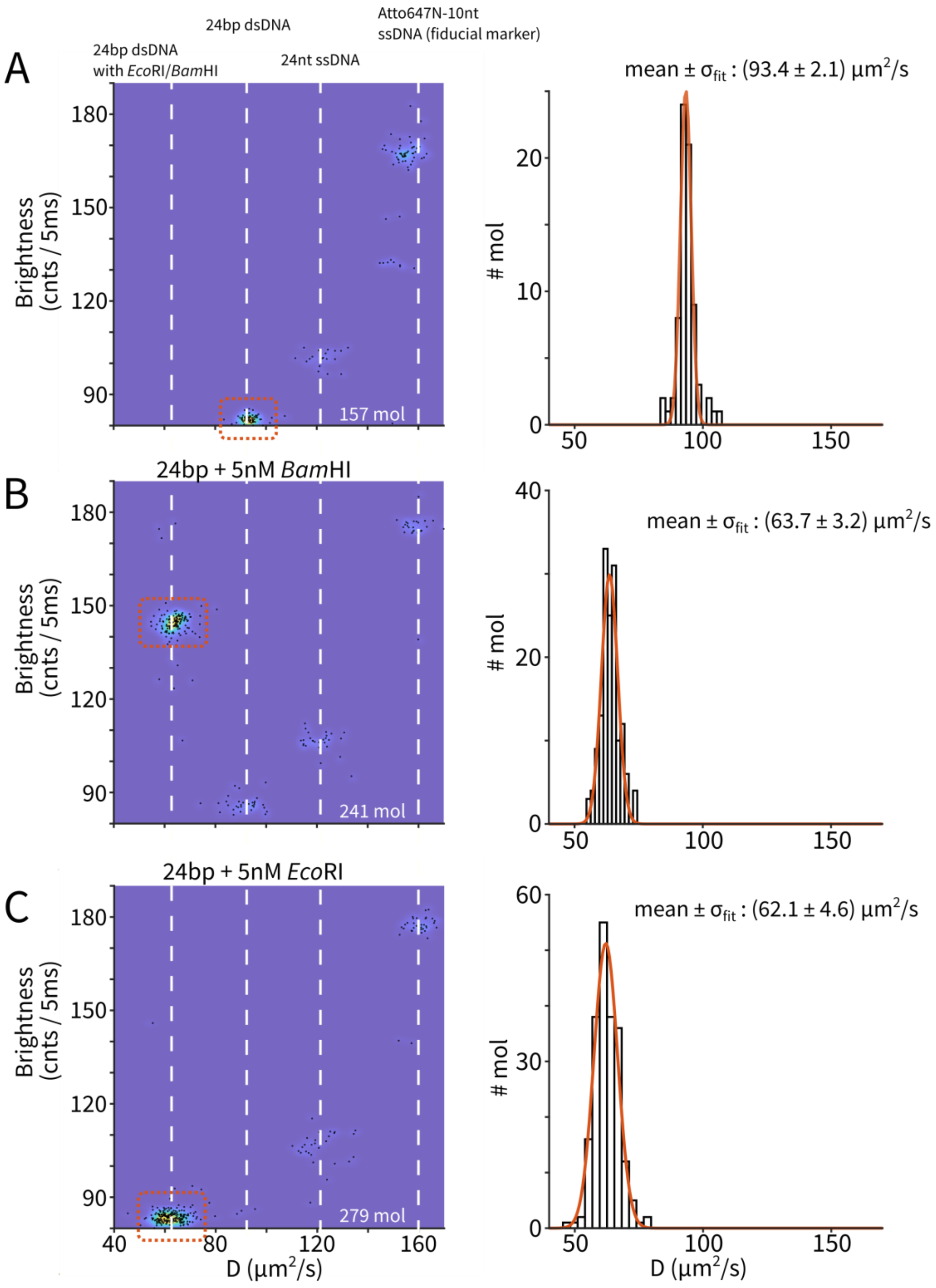


**Figure S6 Quantitative diffusivity measurement of 24bp DNA, 24bp-*Bam*HI and 24bp-*Eco*RI complexes.** See caption of Figure S5 for details.

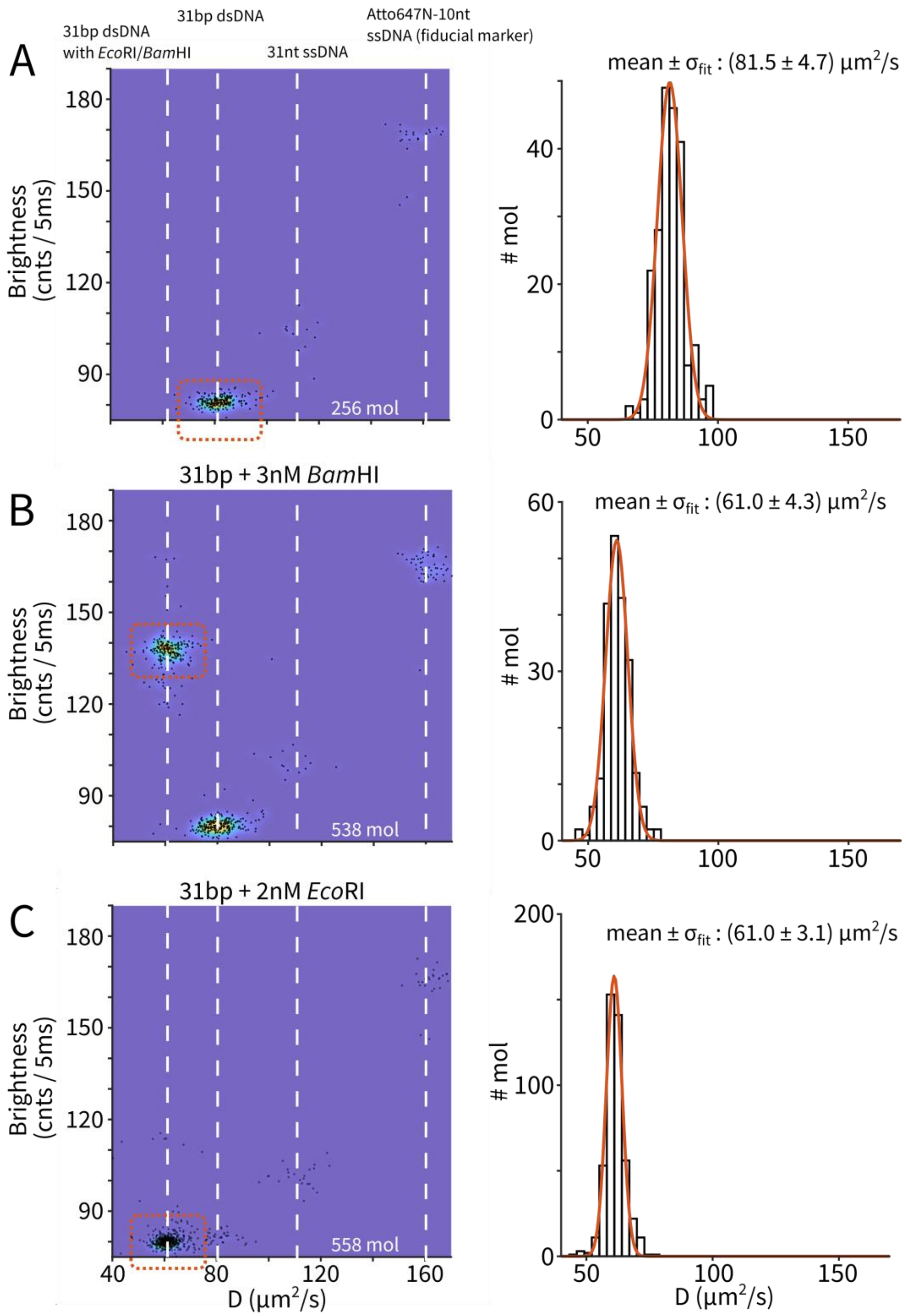


**Figure S7 Quantitative diffusivity measurement of 31bp DNA, 31bp-*Bam*HI and 31bp-*Eco*RI complexes.** See caption of Figure S5 for details.

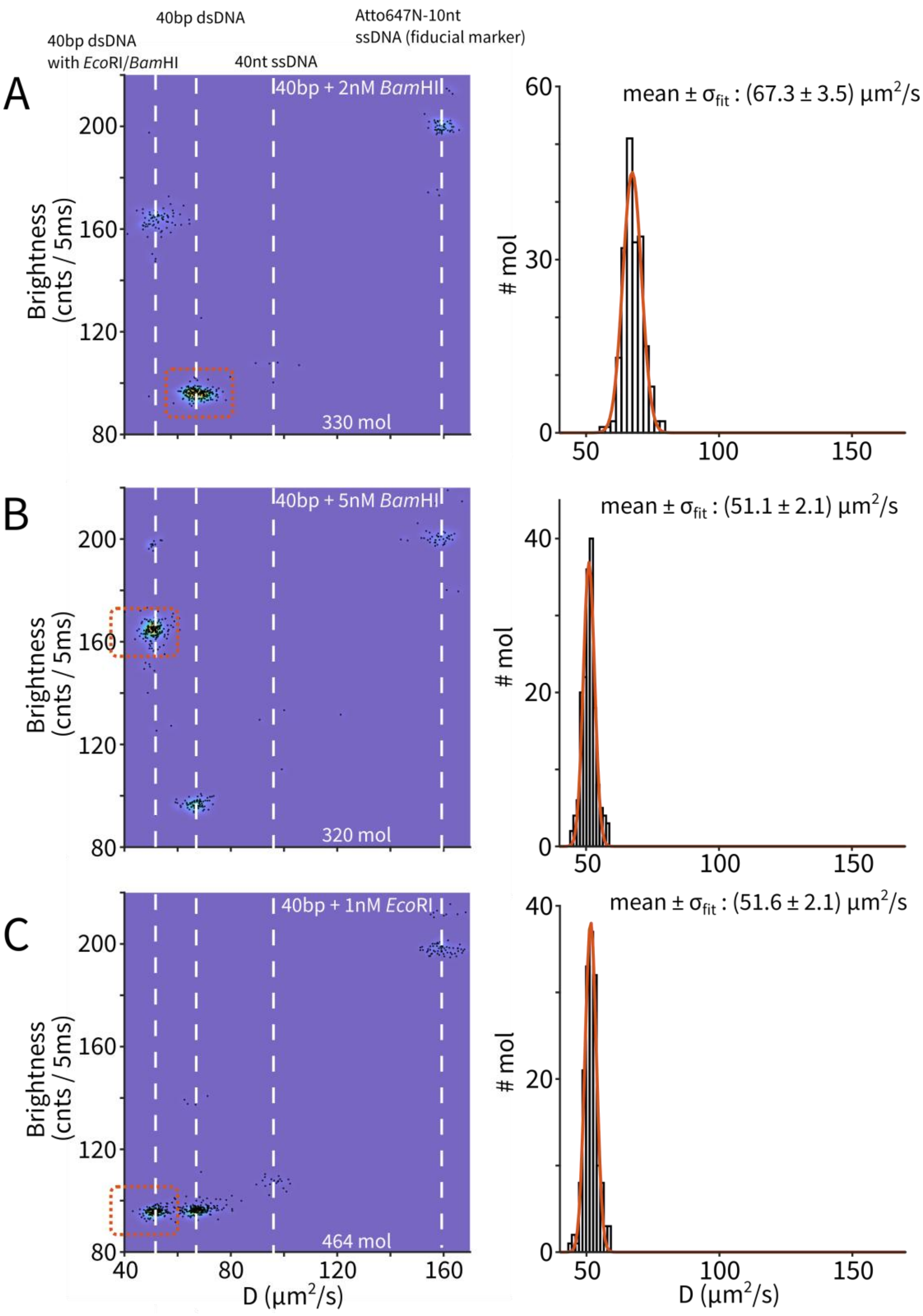


**Figure S8 Quantitative diffusivity measurement of 40bp-*Bam*HI and 40bp-*Eco*RI complexes.**

See caption of Figure S5 for details.

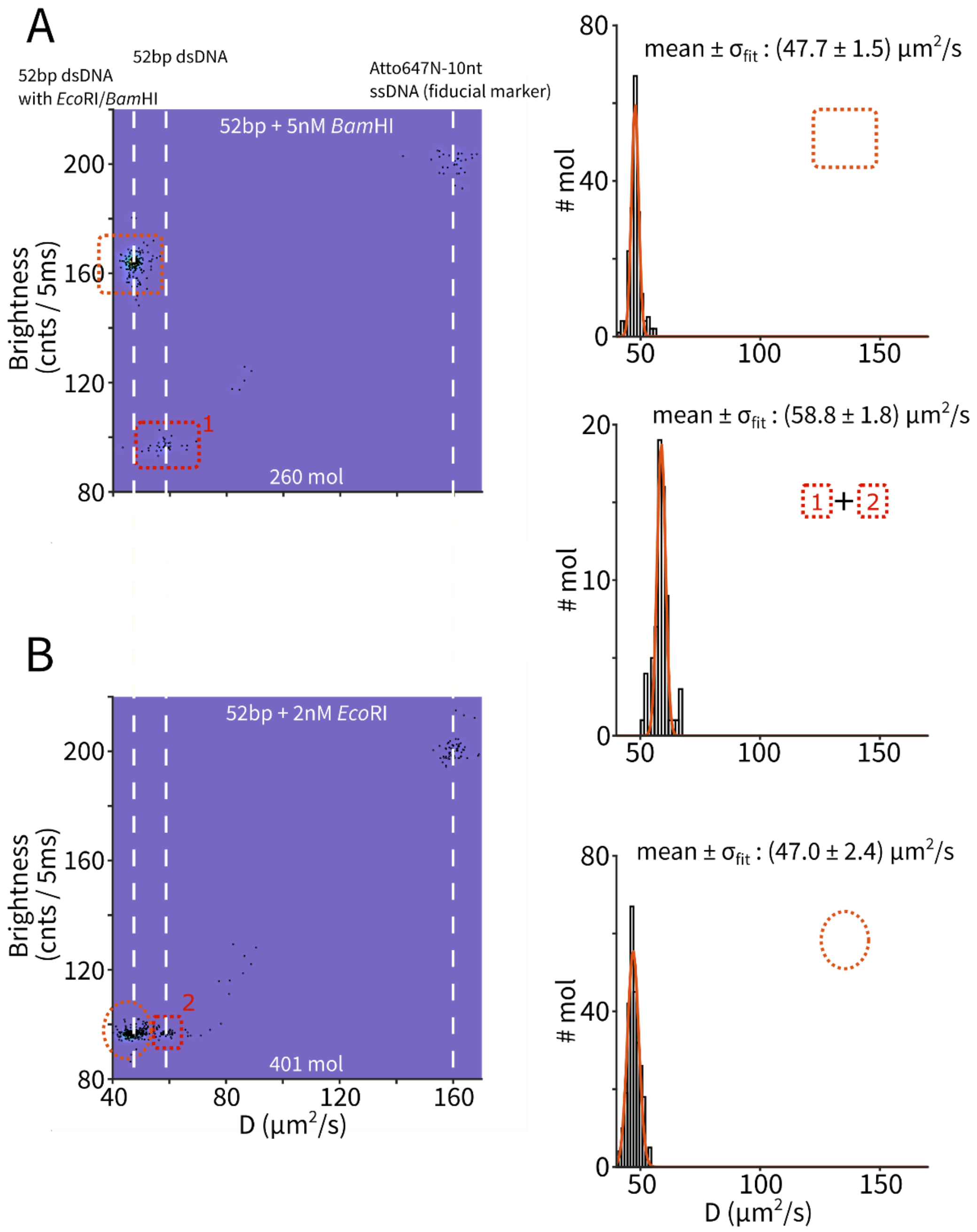


**Figure S9 Quantitative diffusivity measurement of 52bp DNA, 52bp-*Bam*HI and 52bp-*Eco*RI complexes.** See caption of Figure S5 for details. In this experiment, the diffusion coefficient of the free DNA (middle panel on the right) was estimated by pooling the DNA-only populations from panels A and B (red rectangles labeled 1 and 2).

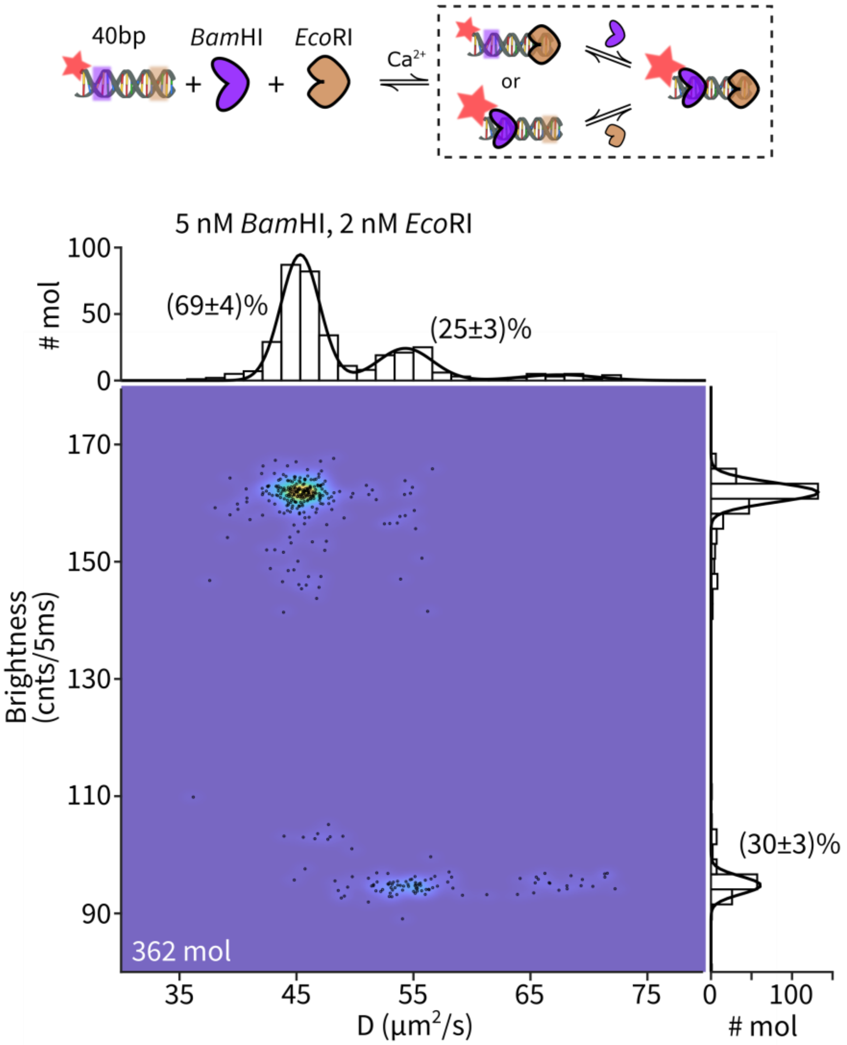


**Figure S10 Combining single-molecule diffusivity contrast and PIFE identifies all four binding configurations between a dual-binding-site DNA and two proteins.** Experimental diffusivity-brightness scatter plot of 362 measured molecules with 5 nM *Bam*HI and 2 nM *Eco*RI. The marginal distribution along the brightness axis was fit with a 2-component Gaussian distribution while the D marginal distribution was fit with a 3-component Gaussian distribution.

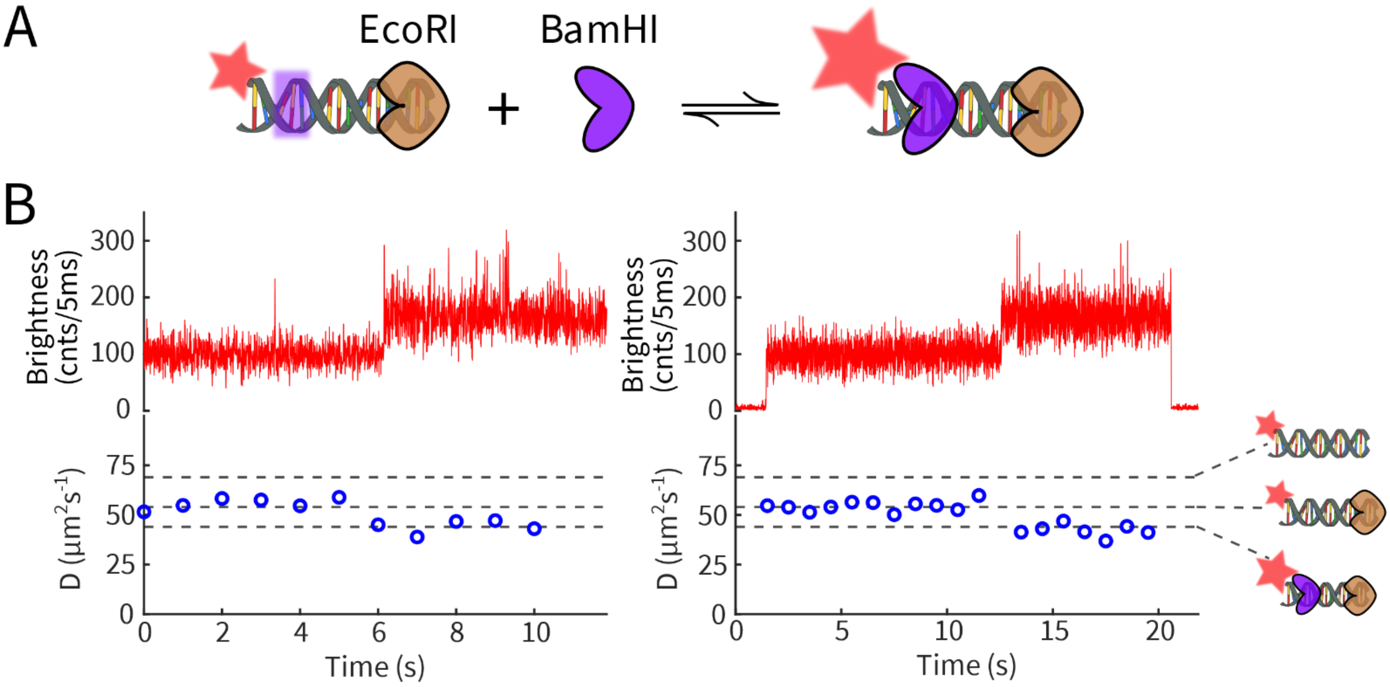


**Figure S11 Sensing dynamic assembly using PIFE and single-molecule diffusometry (a)** Cartoon showing *Bam*HI binding to *Eco*RI-bound 40bp duplex DNA. **(b)** Two example measurements consistent with *Bam*HI binding to DNA-*Eco*RI to form the fully assembled, ternary complex in real-time. Each example shows the brightness (top) and diffusion coefficient (*D*, bottom) over time. *D* is estimated for consecutive 1 second segments of the data. The horizontal dashed lines indicate the expected diffusion coefficient for unbound DNA, DNA-*Eco*RI, and DNA-*Eco*RI-*Bam*HI (cartoons on the right). Note that in these two examples, the intermediate *D* state (~53 $\mu m^2/s$) is assigned as DNA-*Eco*RI complex. DNA-*Bam*HI would have a similar *D* but increased brightness.

**Supplementary Code: extHullRad.py**

Python code that implements the extHullRad pipeline for predicting the hydrodynamic properties of DNA-protein complexes. Instructions to use the code are provided in comments.
