## Supplementary Code for "Probing DNA-protein interactions using single-molecule diffusivity contrast": extHullRad.py

```
#!/usr/bin/env python3
"""
First line is linux related header to make sure python3 is used

extHullRad
Calculates hydrodynamic coefficients for a biomolecular structure. Allows
substitution of the DNA strand for a rod model, and shifting with respect to 
the centroid of the strand.
AUTHOR: Miles Lee, Princeton University
DATE: 2018

Uses code adapted from:
HullRad
AUTHOR: Patrick Fleming
DATE:2017

Fleming, PJ and Fleming, KG "HullRad: Fast Calculations of Folded and Disordered
Protein and Nucleic Acid Hydrogynamic Properties", Biophysical Journal, 2018
DOI: 10.1016/j.bpj.2018.01.002


--------------------------------------------------------------------------------
Requires python 3

Uses PDB file format as input.

Requires qconvex in path. This is a program in the qhull suite of programs.
Qhull may be downloaded from http://www.qhull.org/download.
	For UNIX download "Qhull_2015.2 for Unix"
	tar xzvf qhull-2015-src-7.2.0.tgz
	cd qhull-2015.2/
	make
	sudo make install
	and the path for qconvex will be /usr/local/bin/qconvex

OS X qhull binaries are available from MacPorts package manager. 
	type "sudo port install qhull".
	and the path for qconvex will be /opt/local/bin/qconvex

OS X binaries are also available from Fink package manager.

The default in this script is the UNIX path (/usr/local/bin/qconvex) but you may 
have to change it if you are using OS X.

After installing qhull go to: ### Edit the path to qconvex ### below (~line 658) and
change /usr/local/bin/qconvex to /opt/local/bin/qconvex if that is where qconvex is.

If you are using OS X but are familiar with UNIX and have gcc installed, the UNIX
instructions above work fine on a Mac.

FOR WINDOWS USERS:
There are precompiled versions of the qhull programs available for Windows 7 and up
at http://www.qhull.org/download
Unpack qhull in a directory of choice and ### Edit the path to qconvex ### below (~line 658)
to include the path where your executable was unpacked. 

Output plots a convex hull and molecular structure. Output text looks like:
 
 pdb8rat.ent
 HYDROLASE (NUCLEIC ACID,RNA)            13-AUG-91   8RAT
  M               :        13692     g/mol
  v_bar           :        0.710     mL/g
  Ro(Anhydrous)   :        15.68     Angstroms
  Rg(Anhydrous)   :        14.32     Angstroms
  Dmax            :        45.01     Angstroms
  Axial Ratio     :         1.31
  f/fo            :         1.21
  Dt              :       1.13e-06   cm^2/s
  R(Translation)  :        18.99     Angstroms
  s               :       1.84e-13   sec
  [eta]           :         3.16     cm^3/g
  Dr              :       1.91e+07   s^-1
  R(Rotation)     :        20.36     Angstroms

To write out the unified side chain psuedo-atom model uncomment the two lines
near the bottom labeled, # Write out model (~line 807)

##

----------USAGE-----------
After installing qhull go to: ### Edit the path to qconvex ### below (~line 658)
Before running, please ensure that the input structure is in the same folder 
in .pdb format.
Ensure that the filename, substitute DNA length and positional shift given 
(at ~ line 775) are all correct.
Open PyMOL.
In the PyMOL command line, type:
cd 
Navigate to where this file is saved.
Now type:
run extHullRad.py
PyMOL should now load the hydrodynamic data above the command line, and load
the structure and its respective model convex hulls.


"""

import os
import sys
import math
import string
import shlex, subprocess
import numpy as np

# Imports the compiled graphics object package from pymol to help draw geometric shapes
from pymol.cgo import *
from pymol import cmd

# Radius and rise per base pair of DNA helix in Angstroms
# Use r_dna = 10 and rise_bp = 3.4 for a regular B-DNA double helix
r_dna = 10
rise_bp = 3.4

# Amino Acid mass, volumes,partial specific volumes and unified side chain sphere radius
# Mass is minus water (from SEDNTERP database)
# Volumes from Cohn and Edsall (1943) as corrected by Perkins_EJB_1986 
# vbar is 0.60224(V/M)
#          Mass     Volume     vbar     Radius of equiv. sphere of side chain volume
##ML       Da       A^3    10^-3 m^3/kg   A
AA_data = {
 'ALA': (  71.0939,	87.2,      0.74,	1.852),
 'ARG': ( 156.203,	188.2,     0.73,	3.123),
 'ASN': ( 114.119,	120.1,     0.63,	2.422),
 'ASP': ( 115.104,	115.4,     0.60,	2.356),
 'CYS': ( 103.16,	106.7,     0.62,	2.224),
 'GLN': ( 128.16,	145.1,     0.68,	2.722),
 'GLU': ( 129.131,	140.9,     0.66,	2.676),
 'GLY': (  57.0669,	 60.6,     0.64,	0.000),
 'HIS': ( 137.157,	152.4,     0.67,	2.798),
 'HSE': ( 137.157,	152.4,     0.67,	2.798),
 'HSP': ( 137.157,	152.4,     0.67,	2.798),
 'HSD': ( 137.157,	152.4,     0.67,	2.798),
 'ILE': ( 113.175,	168.9,     0.90,	2.957),
 'LEU': ( 113.175,	168.9,     0.90,	2.957),
 'LYS': ( 128.19,	174.3,     0.82,	3.005),
 'MET': ( 131.215,	163.1,     0.75,	2.903),
 'MSE': ( 178.125,	163.1,     0.75,	2.903),
 'PHE': ( 147.192,	187.9,     0.77,	3.121),
 'PRO': (  97.132,	122.4,     0.76,	2.453),
 'SER': (  87.093,	 91.0,     0.63,	1.936),
 'THR': ( 101.12,	117.4,     0.70,	2.385),
 'TRP': ( 186.229,	228.5,     0.74,	3.422),
 'TYR': ( 163.192,	192.1,     0.71,	3.155),
 'VAL': (  99.148,	141.4,     0.86,	2.682)
}

# From voss_jmb_2005.pdf and nadassy_nar_2001.pdf
# Data for nucleoside, = nucleotide - 79.95 for PO3H
# Phosphate for na lacking terminal phosphates
# vbar is 0.60224(V/M)
#          Mass         Volume  vbar    Radius of equiv. sphere of base volume
##         Da           A^3    10^-3 m^3/kg   A
NA_data = {
 '  A':  ( 267.25,       272.3,  0.614,   0.00),
 '1MA':  ( 281.25,       299.3,  0.641,   0.00),
 'MIA':  ( 383.45,       385.1,  0.605,   0.00),
 '  C':  ( 243.22,       248.1,  0.614,   0.00),
 '5MC':  ( 257.22,       275.1,  0.644,   0.00),
 'OMC':  ( 257.22,       275.1,  0.644,   0.00),
 '  G':  ( 283.24,       279.0,  0.593,   0.00),
 '2MG':  ( 297.27,       306.0,  0.620,   0.00),
 '7MG':  ( 299.27,       306.0,  0.620,   0.00),
 'M2G':  ( 311.27,       333.0,  0.644,   0.00),
 'OMG':  ( 297.27,       306.0,  0.620,   0.00),
 'YYG':  ( 508.54,       500.0,  0.593,   0.00),
 ' YG':  ( 428.55,       427.1,  0.600,   0.00),
 '  G':  ( 283.24,       279.0,  0.593,   0.00),
 '  U':  ( 244.20,       243.9,  0.602,   0.00),
 '5MU':  ( 258.20,       270.9,  0.632,   0.00),
 '4SU':  ( 260.35,       270.5,  0.626,   0.00),
 'H2U':  ( 246.20,       243.9,  0.602,   0.00),
 'PSU':  ( 244.20,       243.9,  0.602,   0.00),
 '  I':  ( 268.23,       273.0,  0.613,   0.00),
 ' DA':  ( 251.25,       271.0,  0.650,   0.00),
 ' DC':  ( 227.22,       246.8,  0.654,   0.00),
 ' DG':  ( 267.25,       277.7,  0.626,   0.00),
 ' DT':  ( 242.23,       264.4,  0.657,   0.00),
 ' DI':  ( 252.22,       271.7,  0.649,   0.00),
 'PO2':  (  62.97,	      52.4,  0.501,   0.00)
}

# From HullRad
def distance(ax,ay,az, bx,by,bz):
    #Euclidean distance between two atoms
    return math.sqrt((ax - bx)**2.0 + (ay - by)**2.0 + (az - bz)**2.0)

# From HullRad
def get_coords(model_array):
    # Get x,y,z coords of CA and CB atoms
    ## Carbon alpha and beta coordinates (backbone of sequence)
    coords = []
    for row in range(len(model_array)):
        coords.append((float(model_array[row][4]), float(model_array[row][5]), float(model_array[row][6])))
    return coords

# Adapted from HullRad. Builds atomic model of protein and DNA seperately.
def protein_model_from_pdb(file, use_new_model):
    # Makes a reduced atom  model of the pdb file
    # This is the model used to make the convex hull
    # For proteins the CB is displaced along the CA-CB vector a distance 
    #   equal to the radius of a sphere equal to the volume of the side chain
    # For nucleic acids only the oxygen and nitrogen atoms are used

    ## When use_new_model is True, this function builds USC model for the amino acid 
    ## if carbon beta is present, and just uses the backbone atoms if the carbon beta
    ## is absent. When use_new_model is False, this function works in the same way as
    ## given in the original HullRad
 
    # Read the PDB file
    data = open(file, 'r').readlines()
 
    # Initialize
    # Initial protein atoms from PDB file
    prot_rec = []
    # Initial nucleic acid atoms from PDB file
    na_rec = []
    # Reduced list of atoms
    atom_rec = []
    struc_name = ' '
    num_MG = 0
    num_MN = 0

    # Get all relevant atoms even if in wrong order
    for line in data:
        # Get name of structure
        if (line[:6] == 'HEADER'):
            struc_name = line.strip()
            print(struc_name[9:])

        elif (line[:4] == 'ATOM'):
            resname = line[17:20]
            if (resname in AA_data) and (line[13:15] == 'N ' or \
              line[13:15] == 'CA' or \
              line[13:15] == 'C ' or \
              line[13:15] == 'O ' or \
              line[13:16] == 'OT1' or \
              line[13:15] == 'CB') and \
              (line[16] != 'B') and \
              (line[:3] != 'END'):
                prot_rec.append(line)
            elif (resname in NA_data) and (line[13:14] == 'N' or \
              line[13:14] == 'O' or line[13:14] == 'P'):
                na_rec.append(line)
        elif (line[:6] == 'HETATM'):
            resname = line[17:20]
            if line[12:14] == 'MG':
                num_MG += 1
            elif line[12:14] == 'MN':
                num_MN += 1
            # Some nucleic acids are in HETATM (!)
            elif (resname in NA_data) and (line[13:14] == 'N' or \
              line[13:14] == 'O' or line[13:14] == 'P'):
                na_rec.append(line)

    # Get only the CB atoms from the protein
    cb_rec = []
    for line in prot_rec:
        if (line[13:15] == 'CB') and \
           (line[0:3] != 'END'):
            cb_rec.append((line))

    # Count the CB atoms and add to atom list in correct order
    num_cb = len(cb_rec)
    line_counter = 0
    cb_counter = 0
    if use_new_model:
        ## Count number of missing CB atoms
        missing_cb = 0
        for line in prot_rec:
            if (line[13:15] == 'N ' or \
              line[13:15] == 'CA' or \
              line[13:15] == 'C ' or \
              line[13:15] == 'O ' or \
              line[13:16] == 'OT1'):
                atom_rec.append((line))
                line_counter += 1
                if (math.fmod(line_counter,4) == 3):
                    ## Before checking for CB, note the index of the residue
                    resIndex = line[22:26]
                if (math.fmod(line_counter,4) == 0 and line[17:20] != 'GLY' \
                     and cb_counter < num_cb):
                    ## Only add CB if it belongs to same residue in the sequence
                    if (cb_rec[cb_counter][22:26] == resIndex):
                        atom_rec.append((cb_rec[cb_counter]))
                        cb_counter +=1
                    else: 
                        ## Otherwise the CB is missing
                        missing_cb += 1
                elif (math.fmod(line_counter,4) == 0 and line[17:20] != 'GLY' \
                     and cb_counter >= num_cb):
                    print(' Error: ATOM entries in pdb file incorrectly ordered')
                    sys.exit()
        
        ## Report the number of missing side chains due to missing CB
        if (missing_cb != 0):
            if (missing_cb == 1):
                print('Caution: there is 1 missing side chain ' +
                      'in the unified side chain model')
            else:
                print('Warning: there are ' + str(missing_cb) + 
                      ' missing side chains in the unified side chain model')

    else:
        for line in prot_rec:
            if (line[13:15] == 'N ' or \
              line[13:15] == 'CA' or \
              line[13:15] == 'C ' or \
              line[13:15] == 'O ' or \
              line[13:16] == 'OT1'):
                atom_rec.append((line))
                line_counter += 1
                if (math.fmod(line_counter,4) == 0 and line[17:20] != 'GLY' \
                     and cb_counter < num_cb):
                    atom_rec.append((cb_rec[cb_counter]))
                    cb_counter +=1
                elif (math.fmod(line_counter,4) == 0 and line[17:20] != 'GLY' \
                     and cb_counter >= num_cb):
                    print(' Found non-gly residue with no CB atom')
                    sys.exit()

    # Put atom info into model array
    num_atoms =  len(atom_rec)
    model_array = [['X' for j in range(7)] for i in range(num_atoms)]
    for row in range(len(model_array)):
        model_array[row][0] = row
        model_array[row][1] = (atom_rec[row][11:16])
        model_array[row][2] = (atom_rec[row][17:20])
        model_array[row][3] = (atom_rec[row][22:26])
        model_array[row][4] = (atom_rec[row][30:38])
        model_array[row][5] = (atom_rec[row][38:46])
        model_array[row][6] = (atom_rec[row][46:54])

    # Make unified side chain as CB
    for row in range(len(model_array)):
        if (model_array[row][1] == '  CA ' and model_array[row][2] != 'GLY' \
             and model_array[row+3][1] == '  CB '):
            rnam = model_array[row][2]
            # CA coords
            ca_x = float(model_array[row][4])
            ca_y = float(model_array[row][5])
            ca_z = float(model_array[row][6])
            # CB coords (third next coord: C,O,CB)
            cb_x = float(model_array[row+3][4])
            cb_y = float(model_array[row+3][5])
            cb_z = float(model_array[row+3][6])
            # CA-CB distance
            dx = cb_x - ca_x
            dy = cb_y - ca_y
            dz = cb_z - ca_z
            ca_cb_dist = distance(ca_x,ca_y,ca_z,cb_x,cb_y,cb_z)
            # Extend CB outward along CA-CB vector
            aa_mass, vol_aa, vbar_aa, sc_rad_aa = AA_data[rnam]            
            new_cb_x = ca_x + (sc_rad_aa * dx/ca_cb_dist)
            new_cb_y = ca_y + (sc_rad_aa * dy/ca_cb_dist)
            new_cb_z = ca_z + (sc_rad_aa * dz/ca_cb_dist)
            model_array[row+3][4] = str(new_cb_x)
            model_array[row+3][5] = str(new_cb_y)
            model_array[row+3][6] = str(new_cb_z)

    # Put nucleic acid atoms into initial array
    num_atoms =  len(na_rec)
    na_array = [['X' for j in range(7)] for i in range(num_atoms)]
    for row in range(len(na_array)):
        na_array[row][0] = row
        na_array[row][1] = (na_rec[row][11:16])
        na_array[row][2] = (na_rec[row][17:20])
        na_array[row][3] = (na_rec[row][22:26])
        na_array[row][4] = (na_rec[row][30:38])
        na_array[row][5] = (na_rec[row][38:46])
        na_array[row][6] = (na_rec[row][46:54])

    return num_MG,num_MN,model_array,na_array

# Finds a line of best fit through the DNA in the sample
def linreg(na_array, n_bp, rise_bp, shift):
    x = []
    y = []
    z = []

    for row in range(len(na_array)): 
        x.append(float(na_array[row][4]))
        y.append(float(na_array[row][5]))
        z.append(float(na_array[row][6]))

    x_array = np.asarray(x)
    y_array = np.asarray(y)
    z_array = np.asarray(z)
    data = np.concatenate((x_array[:, np.newaxis],  y_array[:, np.newaxis], z_array[:, np.newaxis]), axis=1)

    # Finds the center of the rod
    datamean = data.mean(axis=0)

    # Performs a Single Value Decomposition on the centered data.
    u, s, vh = np.linalg.svd(data - datamean)

    # vh[0] contains the first principal component, the direction
    # vector of the 'best fit' line (helical axis)

    # Extends a rod of desired length along the helical axis
    rad = n_bp * rise_bp / 2
    linepts = vh[0] * (shift * rise_bp + np.mgrid[-rad: rad :2j][:, np.newaxis])
    
    # Shift by the mean to move rod back to original position
    linepts += datamean

    return linepts[0], linepts[1]

# This function gives points at the two ends of cylindrical rod 
def CylEnd(pt1, pt2):
    if r_dna == 0:
        return([pt1, pt2])

    vec = np.subtract(pt2, pt1)
    d = dotProduct(vec, pt2)
    
    if vec[0] != 0:
        pt3 = [d/vec[0], 0, 0]
    elif vec[1] != 0: 
        pt3 = [0, d/vec[1], 0]
    else:
        pt3 = [0, 0, d/vec[2]]
    
    # 4 vectors in end of DNA rod
    u1 = np.subtract(pt3, pt2)
    u1 = np.multiply(u1, r_dna/mag(u1))

    u2 = crossProduct(vec, u1)
    u2 = np.multiply(u2, r_dna/mag(u2))

    u3 = np.add(u1, u2)
    u3 = np.multiply(u3, r_dna/mag(u3))

    u4 = np.subtract(u1, u2)
    u4 = np.multiply(u4, r_dna/mag(u4))
    
    ret = []
    
    # with adding and subtracting, we get 8 points on each end cylinder face
    for u in [u1, u2, u3, u4]:
        for p in [pt1, pt2]:
            ret.append(np.add(p, u))
            ret.append(np.subtract(p, u))
    
    return ret 

# Assumes v is 3d vector. mag(v) gives the magnitude |v|
def mag(v):
    return math.sqrt(v[0] ** 2 + v[1] ** 2 + v[2] ** 2)

def crossProduct(v1, v2):
    ret = []
    ret.append(v1[1] * v2[2] - v1[2] * v2[1])
    ret.append(v1[2] * v2[0] - v1[0] * v2[2])
    ret.append(v1[0] * v2[1] - v1[1] * v2[0])
    return ret

def dotProduct(vt1, vt2):
    ret = vt1[0] * vt2[0] + vt1[1] * vt2[1] + vt1[2] * vt2[2]
    return ret

# From HullRad
def write_pdb(model_array, filename):
    # Write out reduced atom model in PDB format for display
    if type(filename) is type(''):
        ff = open(filename, 'wb')
    else:
        ff = filename

    write = ff.write

    pdbfmt = 'ATOM  %5d%3s %3s  %4d    %8.3f%8.3f%8.3f  0.00  0.00\n'
    for i in range(len(model_array)):
        a = model_array[i]
        write(str.encode((pdbfmt % (int(a[0]),a[1],a[2],int(a[3]),float(a[4]),float(a[5]),float(a[6])))))
    write(str.encode('END\n'))

    ff.close()

# Adapted from HullRad
def Sved(num_MG,num_MN,model_array):
    #
    # Main function: Does most things and calls HullData
    #
    # Use Cohn-Edsall eq. to calc protein specific volume
    # vbar_prot = sum(ni*Mi*vbar_aa) / (ni*Mi) where n is number, M is mass
    #
    prot_mol_mass = 0.0
    numerator = 0.0
    NA = 0

    # Radius of Gyration
    # Calc center of mas
    X = 0.0
    Y = 0.0
    Z = 0.0
    tot = 0.0
    for row in range(len(model_array)):
        X = X + (float(model_array[row][4]))
        Y = Y + (float(model_array[row][5]))
        Z = Z + (float(model_array[row][6]))
        tot += 1

    com_x = (X/tot)
    com_y = (Y/tot)
    com_z = (Z/tot)

    Rg2  = 0.0
    for row in range(len(model_array)):
        if (model_array[row][1] != '  P  '):
            Rg2 += ((distance(com_x, com_y, com_z, float(model_array[row][4]),\
                         float(model_array[row][5]), float(model_array[row][6])))**2)
    Rg = math.sqrt(Rg2/tot)

    # Calc masses for protein and nucleic acid
    for row in range(len(model_array)):
        if (model_array[row][1] == '  CA '):
            rnam = model_array[row][2]
            aa_mass, vol_aa, vbar_aa, sc_rad_aa = AA_data[rnam]            
            numerator += (aa_mass * vbar_aa)
            prot_mol_mass += aa_mass
        elif ((model_array[row][1] == "  O4'") or (model_array[row][1] == "  O4*")):
            rnam = model_array[row][2]
            aa_mass, vol_aa, vbar_aa, sc_rad_aa = NA_data[rnam]
            numerator += (aa_mass * vbar_aa)
            prot_mol_mass += aa_mass
            NA = 1
        # Add PO2 because above were nucleosides
        elif (model_array[row][1] == "  P  "):
            numerator += (62.97 * 0.501)
            prot_mol_mass += 62.97

    # Add weight of water for N-term and C-term of protein
    prot_mol_mass += 18.0

    # If nucleic acid add weight of MG and MN
    ion_mass = (num_MG * 24.305) + (num_MN * 54.938)
    prot_mol_mass += ion_mass
     
    # Correct by -0.0025 because above volumes were measured at 25 deg C and want 20 deg C
    #  as per Svedberg, The Ultracentrifuge, 1940, Oxford Univ. Press
    vbar_prot = (numerator / prot_mol_mass) - 0.0025

    # Get convex hull area and volume 
    coords = get_coords(model_array)
    area_hull, vol_hull, Dmax, obj= HullData(coords)

    # Calculate shell volume from variable of shell thickness
    # Hydration shell thickness is empirically determined to be optimal 
    # This expands each hull plane by hydration thickness
    vol_shell_wat = area_hull * 2.8

    # Calculate total volume of hydrated convex hull (including shell waters)
    vol_hyd_hull = vol_hull + vol_shell_wat

    # Estimate axial ratio 
    # Minus 3 Ang because DNA duplex a should be rod length, not diagonal
    #   of hull end vertices (Also necessary to make apoferritin axial ratio = 1)
    ## ML  Seems rather arbitrary 
    a = (Dmax/2.0) - 3.0
    b = math.sqrt((3.0 * vol_hull)/(4.0*math.pi*a))

    # But some very spherical structures may not have the diagonal a full 3 Ang longer.
    # Can't have a < b.
    if a > b: 
        # Translational Shape Factor
        numerator = math.sqrt(1.0 - (b/a)**2)
        denominator = ((b/a)**0.66666667) * math.log((1 + math.sqrt(1.0 - (b/a)**2))/(b/a))
        Ft = numerator/denominator
    else:
        a = b
        Ft = 1.0

    # Weight shape factor
    # Empirically found to work better with expanded volume
    #  (Many combinations tried)
    Ft = math.sqrt(Ft)

    # Axial ratio of prolate ellipsoid of same volume as convex hull
    a_b_ratio = a/b

    # Find radius of sphere of same volume as hydrated convex hull
    factor = 3.0/(4.0 * math.pi)
    Rht = (factor * vol_hyd_hull)**0.333333
    # Include Shape factor to give effective hydrodynamic translational radius
    Rht = Rht * Ft
    # Rht comes as Angstrom
    # need meters in equation below
    Rh_trans = Rht * 1e-8

    # Calculate Svedberg coeff.
    eta = 0.0100194 # poise
    rho = 0.998234 # g/ml water density
    fT = 6.0*math.pi*eta*Rh_trans
    s = prot_mol_mass * (1.0 - (vbar_prot * rho)) / (6.02214e23 * fT)

    # Calculate translational diffusion coeff.
    #R = 8.314e7
    kB = 1.381e-16
    T = 293.15
    Dt = kB*T / fT

    # Calculate fT/fo
    # Vbar is in ml/g, need A^3/Dalton
    vol_prot = prot_mol_mass*vbar_prot/0.60224
    Ro = ((3.0 * vol_prot) / (4.0 * math.pi))**0.333333
    ffo_hyd_P = Rht/Ro

    # Calculate rotational diffusion coeff.
    # Rotation more strongly affected by hydration, Halle & Davidovic, 2003
    # So increase hydration shell thickness
    # Hydration shell thickness empirically found to be optimal
    vol_shell_wat_rot = area_hull * 4.3
    vol_hyd_hull_rot = vol_hull + vol_shell_wat_rot
    Rhr = (factor * vol_hyd_hull_rot)**0.333333

    # Include empirical correction for shape factor to give effective 
    #  hydrodynamic rotational radius
    # This obtained by fitting to DNA duplexes
    Fr = Ft**4.0
    Rhr = Rhr * Fr
    # Rhr comes as Angstrom
    # need meters in equation below
    Rh_rot = Rhr * 1e-8
    fR = 8.0*math.pi*eta*(Rh_rot**3.0)
    Dr = kB*T / fR

    # Calculate intrinsic viscosity
    # From Einstein viscosity equation,
    # [n] = (2.5*6.02214e23*(4*pi*Rht^3)/3)/prot_mol_mass
    #
    # rearrange to
    # 
    int_vis = (10.0 * math.pi * 0.602214 * ((Rht)**3.0)) / (3.0 * prot_mol_mass)

    return s,Dt,Dr,vbar_prot,Rht,ffo_hyd_P,prot_mol_mass,Ro,Rhr,int_vis,a_b_ratio,Ft,Rg,Dmax,obj

# Adapted from HullRad to include extract convex hull surface triangles
def HullData(coords):
    #
    # Uses qconvex to calculate convex hull
    #
    #   coords are coords of reduced atom model
    #

    # Input for qconvex, etc+
    dim = 3
    num_points = len(coords)
    instring = " %d\n " % (dim )
    instring += " %d\n " % (num_points )
    for i in range(len(coords)):
        instring += ' %f %f %f\n' % (coords[i][0],coords[i][1],
                                   coords[i][2])
    instring += '\n'
    
    ### Edit the path to qconvex ###
    qconvexPath = "D:/HullRad/qhull-2015.2/bin/qconvex"

    # Call qconvex with argument, FA (compute total area and volume)
    p = subprocess.Popen([qconvexPath, "FA"],stdin=subprocess.PIPE, stdout=subprocess.PIPE, stderr=subprocess.PIPE)
    (input, output) = (p.stdin, p.stdout)
    input.write(str.encode(instring))
    input.close()
    data = output.read()

    # Area of convex hull
    area_start = data.find(str.encode("area:"))
    area_end = data.find(str.encode("Total volume:"))
    if area_end == -1:
        area_end = data.find(str.encode("Approximate volume:"))
    area_hull = float(data[(area_start + 5) : area_end] )

    # Volume of convex hull
    vol_start = data.find(str.encode("volume:"))
    if vol_start == -1 :
        print(data)
        return 0
    lastStr = data[vol_start + 7 :].split()
    vol_hull = float(lastStr[0])

    # Find max distance between vertices for Dmax calculation
    # Necessary for length of corresponding prolate ellipsoid of revolution
    # Run qconvex with different argument, p (returns vertex coordinates)
    p = subprocess.Popen([qconvexPath, "p"],stdin=subprocess.PIPE, stdout=subprocess.PIPE, stderr=subprocess.PIPE)
    (input, output) = (p.stdin, p.stdout)
    input.write(str.encode(instring))
    input.close()
    data = map(string.split, output.readlines())
    Dmax = 0.0
    for i in range(2,len(data)):
        for j in range(3,len(data)):
            dist = math.sqrt((float(data[i][0])-float(data[j][0]))**2.0 + \
                             (float(data[i][1])-float(data[j][1]))**2.0 + \
                             (float(data[i][2])-float(data[j][2]))**2.0 )
            if dist > Dmax:
                Dmax = dist


    # Calls qconvex to find facets for plotting 
    p = subprocess.Popen([qconvexPath, "i"],stdin=subprocess.PIPE, stdout=subprocess.PIPE, stderr=subprocess.PIPE)
    (input, output) = (p.stdin, p.stdout)
    input.write(str.encode(instring))
    input.close()
    data = map(string.split, output.readlines())
    obj  = []

    # For each facet in the data set draw a facet
    for i in range(1, len(data)):

        # Initialise a face
        obj.extend([
            BEGIN, TRIANGLE_FAN,
            COLOR, 0.5, 0.5, 0.5,
            ALPHA, 0.5,
            ])
        
        # Record vertices
        vx = []
        vy = []
        vz = []
        
        # Collect the vertices
        for j in range(len(data[i])):

            obj.append(
                VERTEX,   )

            x = coords[int(data[i][j])][0]
            y = coords[int(data[i][j])][1]
            z = coords[int(data[i][j])][2]
            
            obj.append(float(coords[int(data[i][j])][0]))
            obj.append(float(coords[int(data[i][j])][1]))
            obj.append(float(coords[int(data[i][j])][2]))
            
            if j < 3:
                vx.append(x)
                vy.append(y)
                vz.append(z)

        nx, ny, nz = normal(vx, vy, vz)

        obj.extend([NORMAL, nx, ny, nz,])
        obj.append(
            END,
            )

    return area_hull, vol_hull, Dmax, obj

# Computes normal vector given first 3 coordinates (a, b, c) in format vx = [ax, bx, cx]
# vy = [ay, by, cy], vz = [az, bz, cz]
def normal(vx, vy, vz):
    u1 = vx[1] - vx[0]
    u2 = vy[1] - vy[0]
    u3 = vz[1] - vz[0]
    v1 = vx[2] - vx[0]
    v2 = vy[2] - vy[0]
    v3 = vz[2] - vz[0]
    n1 = u2 * v3 - u3 * v2
    n2 = u3 * v1 - u1 * v3
    n3 = u1 * v2 - u2 * v1
    mg = -(n1 ** 2 + n2 ** 2 + n3 **2) ** 0.5
    n1 = n1 / mg 
    n2 = n2 / mg
    n3 = n3 / mg 
    return n1, n2, n3

# Use HullRad2.py algorithm for constructing USC atomic model
# Change to False if you want to use the original HullRad
# Setting to False may throw errors for missing beta carbons
use_new_model = True

# name of pdb file 
file = "1eri_dimer.pdb"

# number of base pairs in substituted DNA
n_bp = 40

# How many base pairs along the helical axis the DNA rod should be shifted
# please determine this by examining the locations of the binding site as
# given in the original pdb file and substituted DNA sequence.
# Note that a shift of 0 corresponds to aligning the center of the rods, and
# not the binding site. Also, you may have to change the sign of the shift,
# if the generated DNA is shifted in the opposite direction that you want.
shift = 0

# Make the reduce atom model seperatedly for protein and DNA
num_MG,num_MN,model_array, na_array = protein_model_from_pdb(file, use_new_model)

if na_array == []:
    print("No nucleic acids found to build rod model.")
    sys.exit()

n = len(model_array)

# Find coordinates of points on the end of the DNA rod
endpts = linreg(na_array, n_bp, rise_bp, shift)
pts = CylEnd(endpts[0], endpts[1])

# Add these artifical points to the model array in a processable format 
cnt = 1
for p in pts:
    atm = [cnt,  "  04'", ' DA', '   ' + str(cnt)]
    atm.extend(p)
    model_array.append(atm)
    cnt += 1 

# Write out model
# Uncomment next two lines if you want the cylinder DNA model written out
# basename = os.path.splitext(file)[0]
# write_pdb(model_array, basename + '_model.pdb')

# From HullRad but extracts obj which includes the convex hull faces
# Call the main function
s,Dt,Dr,vbar_prot,Rht,ffo_hyd_P,M,Ro,Rhr,int_vis,a_b_ratio,Ft,Rg,Dmax,obj = \
       Sved(num_MG,num_MN,model_array)


# Change the folowing to True to print HullRad Hydrodynamics data calculation.
giveHydrodynamicData = True
if (giveHydrodynamicData):
    # Print coefficients to screen
    # Many are commented out since model is inaccurate for those attributes.
    # print('  M               :       %6.0f     g/mol' % (M))
    # print('  v_bar           :       %6.3f     mL/g'  % (vbar_prot))
    # print('  Ro(Anhydrous)   :       %6.2f     Angstroms' % (Ro))
    # print('  Rg(Anhydrous)   :       %6.2f     Angstroms' % (Rg))
    print('  Dmax            :       %6.2f     Angstroms' % (Dmax))
    print('  Axial Ratio     :       %6.2f' % (a_b_ratio))
    print('  f/fo            :       %6.2f'  % (ffo_hyd_P))
    print('  Dt              :       %6.2e   cm^2/s' % (Dt))
    print('  R(Translation)  :       %6.2f     Angstroms' % (Rht))
    # print('  s               :       %6.2e   sec' % (s))
    # print('  [eta]           :       %6.2f     cm^3/g' % (int_vis))
    # if a_b_ratio > 2.63:
    #     print(' ')
    #     print('  Caution. Axial ratio too large for accurate prediction')
    #     print('  of following rotational properties by HullRad.')

    # print('  Dr              :       %6.2e   s^-1' % (Dr))
    # print('  R(Rotation)     :       %6.2f     Angstroms' % (Rhr))

# print(obj)
cmd.load(file)
cmd.load_cgo(obj, file.split('.')[0] + '_ModelHull')
cmd.set("light", [2500, 2500, 1000])
cmd.set("two_sided_lighting", 'on')
```
